# Spatiotemporal Dynamics of Intra-tumoral Dependence on NEK2-EZH2 Signaling in Glioblastoma Cancer Progression

**DOI:** 10.1101/2020.12.01.405696

**Authors:** Jia Wang, Marat S Pavliukov, Daisuke Yamashita, Peng Cheng, Zhuo Zhang, Sung-Hak Kim, Mayu A Nakano, Wanfu Xie, Dongquan Chen, Brendan Frett, Wen-hao Hu, Yong Jae Shin, Yeri Lee, Violaine Goidts, Do-Hyun Nam, Hong-yu Li, Ichiro Nakano

**Author notes:** To whom correspondence should be addressed: Ichiro Nakano, MD, PhD. Equal contribution to this study (Jia Wang, Marat S Pavliukov).

## Abstract

The highly lethal brain cancer glioblastoma undergoes dynamic changes in molecular profile and cellular phenotype throughout tumor core establishment and in primary-to-recurrent tumor progression. These dynamic changes allow glioblastoma tumors to escape from multimodal therapies, resulting in patient lethality. Here, we identified the emergence of dependence on NEK2-mediated EZH2 signaling, specifically in therapy-resistant tumor core-located glioblastoma cells. In patient-derived glioblastoma core models, NEK2 was required for *in vivo* tumor initiation, propagation, and radio-resistance. Mechanistically, in glioblastoma core cells, NEK2 binds with EZH2 to prevent its proteasome-mediated degradation in a kinase-dependent manner. Clinically, NEK2 expression is elevated in recurrent tumors after therapeutic failure as opposed to their matched primary untreated cases, and its high expression is indicative of worse prognosis. For therapeutic development, we designed a novel NEK2 kinase inhibitor CMP3a, which effectively attenuated growth of murine glioblastoma models and exhibited a synergistic effect with radiation therapy. Collectively, the emerging NEK2-EZH2 signaling axis is critical in glioblastoma, particularly within the tumor core, and the small molecule inhibitor CMP3a for NEK2 is a potential novel therapeutic agent for glioblastoma.

## Introduction

The devastating intraparenchymal brain tumor glioblastoma presents high intra-tumoral heterogeneity, an obstacle characteristic of various therapy-refractory cancers(*1*). Cellular components of individual tumors continuously undergo dynamic changes both spatially and longitudinally(*2, 3*). As a result, single tumors establish substantial molecular and phenotypic differences between the tumor edge and core (see Supplementary Information for our definition of tumor edge and core), as well as between primary and recurrent tumors(*4*).

The identification of the cell of origin of glioblastoma tumors may not be attainable, at least clinically(*5*). Recent studies, however, have identified some ancestor-like tumor cell subpopulations located at the tumor edge of patient glioblastomas, which presumably give rise to core-located cells later in the tumor development process(*4, 6*). This intra-tumoral Edge-to-Core (E-to-C) progression forces the entire tumor to change phenotypically to adapt to harsh conditions(*4*). Tumor edge-located cells are supported by the rich microenvironment by signals from various neighboring somatic cells (e.g. neuronal BDNF, astrocytic CD38, and vascular endothelial cell-derived endocan)(*7–10*). In contrast, tumor core cells suffer from limited nutrients, driving energy-saving survival competition and growth(*11*).

In developed countries, the initial formation of primary glioblastoma tumors is not the primary cause of patient mortality(*12*). Instead, affected patients die due to the inevitable recurrence of the tumor, following the failure of multi-modal therapies(*12*). These recurrent tumors are predominantly (>80%) found adjacent to the surgical removal cavity (< 2cm)(*13, 14*). Local recurrences, unlike distant ones, are genetically similar to the original primary tumors, suggesting a continuous progression along the primary-to-recurrent (P-to-R) axis(*15*). Analysis of a matched primary and recurrent patient cohort has shown that emergence of the E-to-C molecular signature in recurrence is strongly associated with worse patient prognosis(*4*). The majority of recurrent tumors develops therapy-resistant core lesions through either pre-existing (intrinsic) or acquired (*de novo*) mechanisms(*16–20*). However, molecular determinants for recurrent tumor core development through the E-to-C progression mechanism remain largely unknown.

The epigenetic traits in tumor cells persistently mark and sustain their phenotypes regardless of tumor microenvironment and inter-cellular crosstalk(*21*). As a self-reinforcing mechanism, histone methylation is maintained consistently, copied from mother to daughter cells(*21*). As a master epigenetic regulator, EZH2 suppresses gene transcriptional machinery primarily by catalyzing the trimethylation of histone H3 at lysine 27 (H3K27me3). EZH2 expression is elevated in various human cancer types including glioblastoma, and is associated with tumor malignancy and poor patient outcome(*22–24*). In regards to the intra-tumoral molecular heterogeneity of glioblastoma, increased dependence of peripheral core cells on EZH2 was recently reported, although the comparison was against necrotic tissues inside the enhancing core lesion, both of which are surgically resectable(*11*). In primary (treatment-naive) glioblastoma cells, EZH2-mediated tumor-initiating potential is governed by the oncogenic serine/threonine kinase MELK in combination with multiple cancer-associated transcription factors (e.g. FOXM1, c-JUN, STAT3)(*24–26*). However, a recent study suggests that MELK is likely a functionally-replaceable kinase, at least within several non-brain cancers, although the molecular mechanism for this “replacement” remains undetermined(*27*). To date, therapeutic strategies to target EZH2, or critical regulator(s) for EZH2 signaling, have yet to be achieved clinically.

In this study, we began by investigating molecular persistence and emerging changes in our patient-derived glioblastoma models that develop therapy resistance through therapy-related pressure.

## Results

### EZH2 expression is preferentially localized within the tumor core as opposed to edge

First, we investigated the expression of EZH2 in the tumor core and edge within single glioblastomas *via* immunohistochemistry (IHC). In the exploratory 53 unmatched glioma cohort (44 patient tumor core tissues of varying grades of glioma and 9 tumor edge lesions harboring scattered infiltrative glioblastoma cells), EZH2 expression was markedly elevated in the core tissues compared to the edge (Fig. 1A). IHC with the validation cohort (9 tumor core- and edge-matched glioblastoma cases) confirmed the relatively exclusive expression of EZH2 within the tumor core lesions (Fig. 1B-1C). The publicly available Ivy GBM GAP database supported this data; *EZH2* displayed higher expression patterns within the tumor core-related sublesions, most prominently in cellular tumor (peripheral core lesion) and the least in the leading edge (Fig. 1D). Because the proportion of tumor cells at the core and edge is substantially different (80-90% vs. 5-10%, respectively), we next verified whether this expression difference is in fact within tumor cells and not solely resulted from the lower fraction of tumor cells at the edge. With MRI-guided intra-operative isolation of glioblastoma core and edge tissues by supra-total resection, we established and used tumor core- and edge-derived glioma sphere models and their derivative tumors in mouse brains (Fig. 1E, Fig. S1A-S1C)(*3, 4, 10, 28, 29*). RNA-sequencing (RNA-seq) with these models revealed that *EZH2* expression was significantly higher in the tumor core-derived spheres (n = 3) in comparison to their edge counterparts (n = 2), or normal neural progenitors derived from a human fetal brain (n = 1) (Fig. 1F). Consistent with these *in vitro* data, *EZH2* expression in mouse tumors established from three glioblastoma patients (g0573, g1053, and g1051) was significantly elevated in the tumor core-derived tumors than those from edge lesions (Fig. 1G). We then investigated the efficacy of two EZH2 inhibitors Tazemostat and CPI-1205 on the tumor core- and edge-derived spheres. *In vitro* cell growth assay showed that tumor-core derived spheres were more sensitive to both Tazemostat and CPI-1205 treatment compared to edge-derived spheres (Fig. 1H-1I, Fig. S2A). Moreover, systemic treatment of mice bearing tumor core- or edge-derived sphere tumors with Tazemostat noticeably diminished Ki-67^+^ proliferating tumor cells along with EZH2 expression (Fig. S2B, Fig. 1J). This reduction of tumor cell proliferation was considered, at least in part, due to the microglial shift from anti-tumor M1 phenotype to a pro-tumor M2 one (Fig. S2C-S2D). Taken together, these data suggest that EZH2 is significantly enriched, and functionally operational, in the tumor core-located glioblastoma cells, compared to those at the edge.

**Fig. 1.**
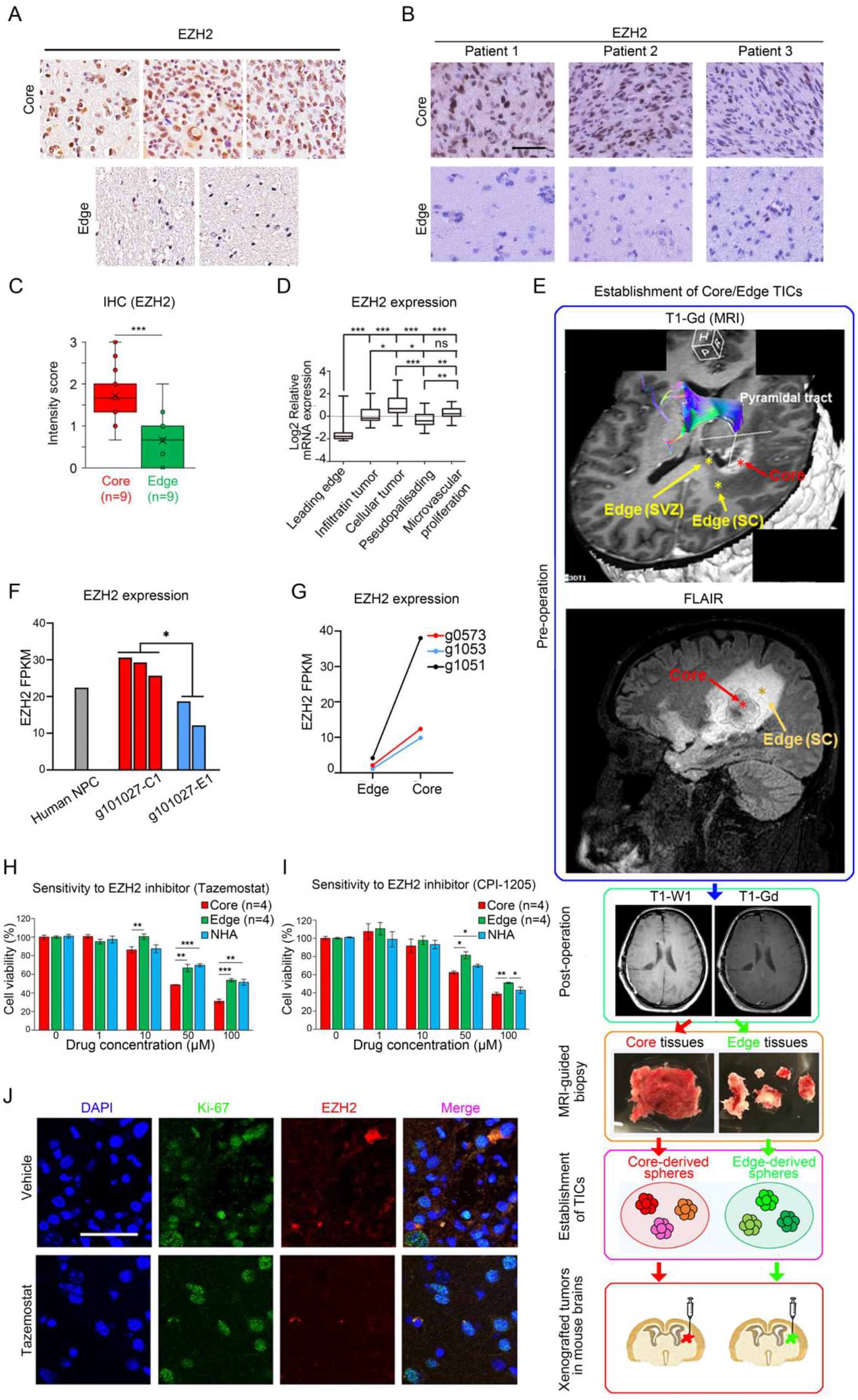
Expression of EZH2 and NEK2 are elevated within the tumor core-located glioblastoma cells and promotes tumor recurrence. (A) Representative immunohistochemical images indicated that EZH2 was significantly enriched in tumor core of glioblastoma compared to the edge area. (B) Representative immunohistochemical images of core and its edge counterparts showed that EZH2 was highly expressed in tumor core (upper panel) compared to tumor edge (lower panel). (C) IHC intensity score analysis indicated that EZH2 expression was remarkably elevated in core area of glioblastoma compared with the edge area (\*\*\**P* < 0.001, with *t* test). (D) Relative mRNA expression for *EZH2* in different region of tumor from Ivy database showed that *EZH2* was enriched in the core-related sublesions (cellular tumor and pseudopalisading) compared to the edge-related sublesions (leading edge) (ns *P* > 0.05, \**P* < 0.05, \*\**P* < 0.01, with one-way ANOVA followed by Dunnett’s *t* tests). (E) Isolation of tumor core- or edge- derived TICs *via* MRI-guided resection of glioblastoma and establishment intracranial mice tumor model. (F) RNA-seq based expression analysis showed that *EZH2* expression was increased in tumor core-derived spheres (n = 3) compared to the edge counterparts and human fetal brain (\**P* < 0. 05, with *t* test). (G) RNA-seq based expression analysis indicated that *EZH2* was highly expressed in the derivative mouse tumors established from the core area of three glioblastoma patients (g0573, g1053, and g1051) compared to their edge counterparts. (H) *In vitro* cell viability assay showed that TICs derived from tumor core were more sensitive to Tazemostat compared to its counterpart derived from the tumor edge, NHA cells were used as a negative control (\*\**P* < 0.01, \*\*\**P* < 0.001, with one-way ANOVA followed by Dunnett’s *t* tests). (I) *In vitro* cell viability assay showed that TICs derived from tumor core were more sensitive to CPI-1205 compared to its counterpart derived from the tumor edge, NHA cells were used as a negative control (\**P* < 0.05, \*\**P* < 0.01, with one-way ANOVA followed by Dunnett’s *t* tests). (J) Immunocytochemistry analysis indicated that Ki67 and EZH2 expression was significantly reduced in mice brain tumor using tumor core-derived TICs then followed by Tazemostat oral treatment. DAPI was used for nuclear staining.

### The serine/threonine kinase NEK2 is highly expressed in glioblastoma core and is elevated during P-to-R progression

The MELK inhibitor OTS167 has been tested in two clinical trials for multiple solid cancers without obvious favorable outcome (https://www.cancer.gov/about-cancer/treatment/clinical-trials/intervention/melk-inhibitor-ots167). Thus, we raised the speculation that cancer cells that underwent long-term OTS167 treatment establish resistance through reactive dependence on the “replaced” kinase. Short-term treatment of our patient-derived glioma sphere lines, all of which were derived from tumor core (n = 7), as well as the conventional glioblastoma cell line U87, resulted in potent effects on *in vitro* growth with the average of the IC50 of 15 nM (Fig. S3A). Likewise, *in vivo* short-term efficacy on U87-derived mouse xenograft models exhibited decreased tumor burden without influencing body weight of the animals (Fig. S3B-S3C). Despite these results, four patient-derived glioma sphere lines treated with the IC50 doses of OTS167 for 4 weeks evoked resistance to both radiation and temozolomide (TMZ) - the current first-line post-surgical therapeutic modalities for glioblastoma (Fig. 2A-2B). Strikingly, these OTS167 long-term treated tumor cells acquired resistance to OTS167 while retained sensitivity to EZH2 inhibition *via* GSK343 treatment (Fig. 2C, Fig. S3D-S3L). These data raise the possibility that some undetermined protein kinase may replace the role of MELK and overcome the effect of OTS167 long-term treatment to subsequently sustain EZH2 activation, thereby retaining, and even promoting glioblastoma’s therapy resistance. Among the candidate kinases, three data sets led us to focus on the poorly characterized serine/threonine kinase NEK2. First, our cDNA microarray data showed that *NEK2* is the most upregulated kinase-encoding gene in tumor core-derived spheres in comparison to normal astrocytes (Fig. S4A)(*16*). Second, hierarchical bi-clustering in 3 previously published GEO databases which were related to glioblastoma (TCGA), TMZ resistance (GSE68029) and radio-resistance (GSE56937) identified *NEK2* as one of 6 kinase-encoding genes significantly correlated with all these three phenotypes, together with *AURKB*, *CDK6*, *CDKN3*, *TRIB2* and *CDK2* (Fig. S4B). Third and most importantly, OTS167-treated U87-derived mouse tumors markedly elevated NEK2 expression, determined by western blotting analysis (Fig. 2D). Regarding intra-tumoral localization, IHC staining of our glioblastoma cohort showed that NEK2 expression was significantly enriched in the core area of glioblastoma compared to the edge (Fig. 2E). Additionally, Germany immunohistochemistry scores (GIS) exhibited that NEK2 was highly expressed in approximately half of the glioblastoma core tissues (25 out of 44) but not in the majority of the tumor edge lesions (1 out of 6) (Fig. 2F). Similar results were observed with the Ivy GAP database, which showed *NEK2* expression as remarkably elevated in tumor core-related area of glioblastoma (Fig. 2G). Similar to the results for EZH2 (Fig. 1F-1G), RNA-seq data for *NEK2* with the glioblastoma core- and edge-derived sphere cultures and their derivative tumors showed that NEK2 mRNA was preferentially expressed in the glioblastoma cells derived from tumor core both *in vitro* and *in vivo* (Fig. 2H-2I). We then compared the expression alternations of kinase-encoding genes between core- and edge-derivative glioblastomas. The results indicated that *NEK2* was one of the top-ranking genes which were enriched in core-derived glioblastoma among the 563 kinase-encoding genes (Fig. 2J). The Human Protein Atlas (THPA) that contained the protein expression profiles with various human cancer types including glioma, lung, colorectal, and breast cancers showed that the NEK2 expression displayed a strong association with EZH2 expression (Fig. 2K, Fig. S4C). Similar to *EZH2*, Rembrandt database demonstrated that *NEK2* mRNA expression in glioblastoma was significantly higher than normal brain and other lower grade glioma groups (Fig. 2L). Regarding the correlation of *NEK2* mRNA expression to post-surgical patient outcome, patients with higher *NEK2* expression demonstrate significantly shorter survival compared to those in the intermediate or low expression groups (Fig. 2M). In addition, another probe for *NEK2* in the Rembrandt database yielded consistent results (Fig. S4D-S4E). As for the protein expression, the NEK2^High^ group of the IHC data had significantly shorter patient survival compared to the NEK2^Low^ group (Fig. 2M). Correlation analysis indicated that NEK2 and EZH2 expression was closely related in these samples (Fig. 2N). Given the substantial elevation of EZH2 expression in recurrent tumors compared to the primary cases in matched glioblastoma patients(*30*), we performed IHC for NEK2 with 5 matched glioblastoma core tissues from the initial surgery with untreated tumors and the second surgery with recurrent local lesions after radiation and TMZ chemotherapy. Although the staining intensities for NEK2 varied among the tested samples from the initial surgeries, we observed an overall trend of elevation of NEK2 in recurrent tumors in comparison to the matched initial tumors (4 out of 5) (Fig. 2P-2Q). Consistent with this data, the proportions of NEK2+ tumor cells within tumors were generally increased in these recurrent tumors (Fig. 2R). Collectively, similar to EZH2EK2 is preferentially expressed by tumor core-located glioblastoma cells and is likely upregulated in the EZH2-dependent, therapy-resistant glioblastoma cells.

**Fig. 2.**
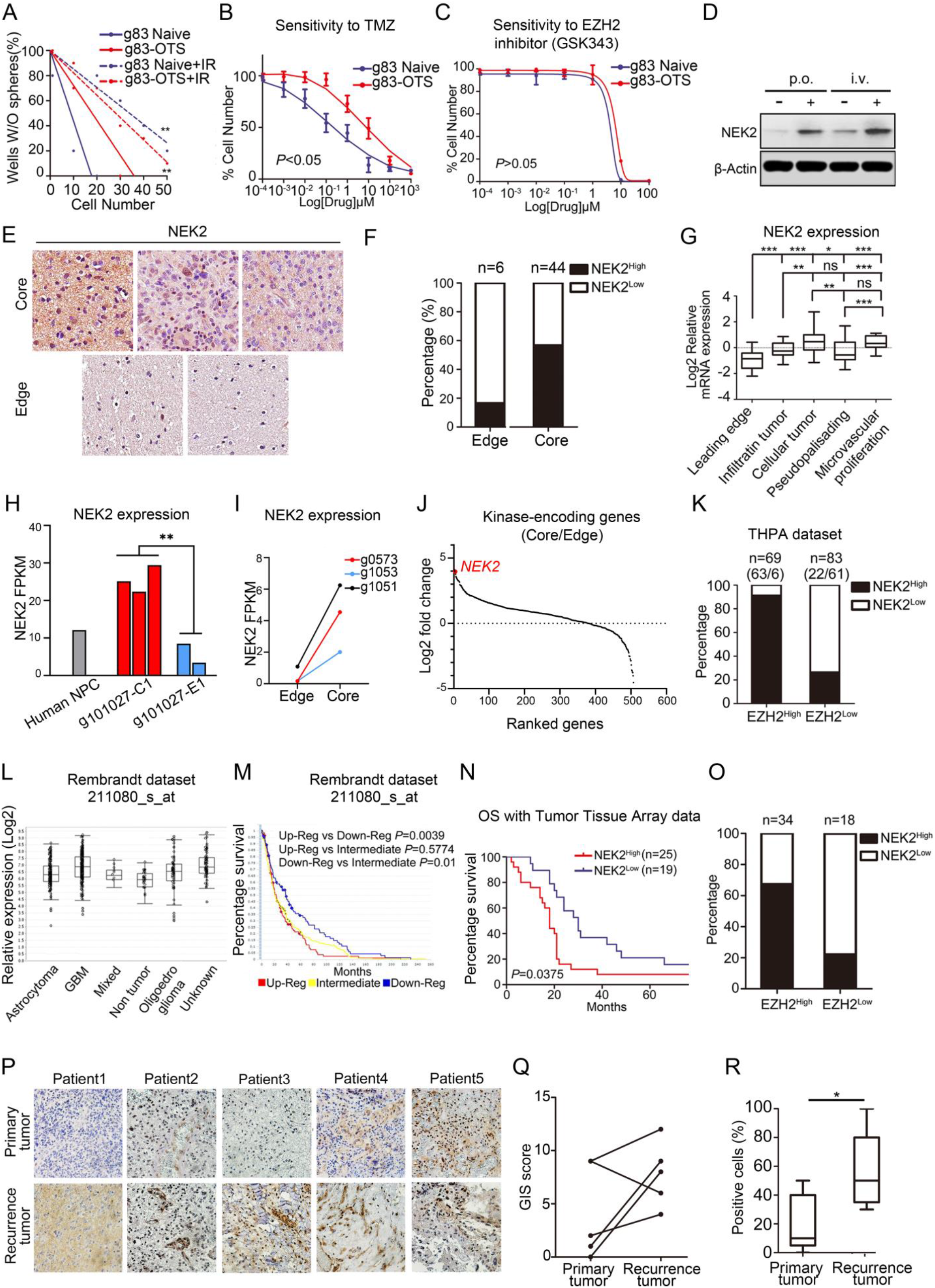
NEK2 is correlated with EZH2 and is a clinically-relevant molecular target in glioblastoma. (A) *In vitro* clonogenicity assay by limiting dilution neuro sphere formation indicated that OTS167 resistant glioma spheres were more resistant to radio-therapy compared with naïve population (\*\**P* < 0.01, with ELDA analysis). (B) *In vitro* cell viability assay showed that OTS167 resistant g83 spheres acquired chemo-resistance to TMZ compared with the naïve cells (*P* < 0.05, with one-way ANOVA). (C) *In vitro* cell viability assay indicated OTS167 resistant population was still sensitive to EZH2 inhibitor GSK343 in g83 spheres (*P* > 0.05, with one-way ANOVA). (D) Western blotting analysis indicated NEK2 expression was increased in U87 sub-Q xenograft mouse model after OTS167 treatment. β-Actin served as a loading control. (E) Representative immunohistochemical images indicated that NEK2 was significantly enriched in tumor core of glioblastoma compared to the edge area. (F) Analysis of NEK2 expression in tumor core and tumor edge of glioma showed that NEK2 was highly expressed in glioma samples (n = 6 in tumor edge, n = 44 in tumor core). (G) Relative mRNA expression for *NEK2* in different region of tumor from Ivy database showed that *NEK2* was enriched in the core-related sublesions (cellular tumor and pseudopalisading) compared to the edge-related sublesions (leading edge) (ns *P* > 0.05, \*\**P* < 0.01, \*\*\**P* < 0.001, with one-way ANOVA followed by Dunnett’s *t* tests). (H) RNA-seq based expression analysis showed that *NEK2* expression was increased in tumor core-derived spheres (n = 3) compared to the edge counterparts and human fetal brain (\*\**P* < 0.01, with *t* test). (I) RNA-seq based expression analysis indicated that *NEK2* was highly expressed in the derivative mouse tumors established from the core area of three glioblastoma patients (g0573, g1053, and g1051) compared to their edge counterparts. (J) RNA-seq based expression analysis indicated that NEK2 was the most up-regulated kinase-encoding genes (ranked fifth among all 563 genes) in core of glioblastoma compared to the edge tumor. (K) NEK2 displayed the strong association with EZH2 expression at protein level in the Human protein Atlas dataset (n = 69 for EZH2^High^ samples and n = 83 for EZH2^Low^ samples). (L) Analysis of the Rembrandt data indicated that *NEK2* expression was elevated in glioblastoma samples (probe set: 211080_s_at). (M) Analysis of the Rembrandt data indicated the inverted correlation between *NEK2* mRNA expression and post-surgical survival of glioma patients (*P* = 0.0071, n = 117 for *NEK2* up-regulated group, n = 314 for *NEK2* intermediate group, n = 110 for *NEK2* down-regulated group, probe set: 211080_s_at). (N) Kaplan-Meier analysis showed significantly negative correlation between NEK2 expression and overall survival in 44 glioma patients (NEK2^High^ samples versus NEK2^Low^ samples, *P* = 0.0375, with log-rank test). (O) Analysis of NEK2 expression in EZH2^High^ and EZH2^Low^ glioma samples (n = 34 for EZH2^High^ samples, n = 18 for EZH2^Low^ samples). Results indicated that NEK2 expression showed significant correlation with EZH2. (P) Immunohistochemically images showed that NEK2 expression was elevated recurrent tumors compared to the primary untreated tumors from 5 matched glioblastoma cases. (Q) NEK2 expression was increased in recurrent tumor compared to the primary counterparts among 5 matched glioblastoma samples. GIS was used to evaluate the expression level of NEK2. (R) NEK2 was highly expressed in recurrent glioblastoma compared to the primary counterparts and presented as percentage of positive stained cells (\**P* < 0.05, with *t* test).

### NEK2 is expressed in radiotherapy-induced tumor-initiating cellular subset in glioblastoma

Next, we investigated whether NEK2 is preferentially expressed in tumor-initiating subpopulation (tumor-initiating cell: TIC) within glioblastoma tumors. We examined the difference of NEK2 expression in the tumor core-derived g528 spheres under two different conditions: serum-free medium with bFGF and EGF (as TIC-enriching condition) and medium with 10% FBS (as differentiation-inducing condition). ICC exhibited that under the TIC-enriching condition, g528 spheres are highly immunoreactive to NEK2 with no detectable expression of a glial differentiation marker GFAP, while these two protein expression patterns were reversed under the differentiation condition (Fig. 3A). Western blotting shows that NEK2 expression is dramatically decreased in g84 and g528 spheres (both of which are derived from tumor core) after treated by serum for 14 days (Fig. 3B). Overall, among 8 patient glioblastoma core-derived sphere lines together with normal human astrocytes, we observed a trend of elevated NEK2 expression in more clonogenic (neuro) sphere-forming lines than others (Fig. S5). *In vitro* radiation to glioma spheres can, at least partially, mimic the glioblastoma core progression by promoting TICs’ therapy resistance, accompanied by their marker shift from CD133 to CD109/Aldehyde Dehydrogenase (ALDH) (*6, 31*). Among 668 kinase-encoding genes, transcriptome microarray data showed that *NEK2* was among the top 13 genes upregulated during the CD133-to-CD109 conversion in 2 glioma sphere samples (gAC17 and g83) (Fig. 3C). qRT-PCR and western blotting confirmed that both NEK2 and EZH2 were similarly elevated in g528 glioma spheres following radiation, which was accompanied by the elevation of CD109 and the decline of CD133 (Fig. 3D-3E). Blocking this CD133-to-CD109 conversion by shRNA for CD109 resulted in a significant reduction of NEK2 expression in g267 glioma spheres (Fig. 3F). When g83 sphere cells was separated into ALDH^High^ TICs and ALDH^Low^ non-TICs by cell sorting, NEK2 was almost exclusively expressed in the ALDH^High^ subpopulation (Fig. 3G). Clinically, expression of both *NEK2* and *EZH2* in 249 glioblastoma samples extracted from the Chinese Glioma Genome Atlas (CGGA) was significantly higher in CD109^High^ samples compared to CD109^Low^ ones (Fig. 3H-3I). Collectively, these data suggested that the NEK2 expression is highly restricted in TICs, particularly radiation-induced ones, in glioblastoma.

### Eliminating NEK2 leads to eradication of glioblastoma-initiating cells in mouse brains

To investigate the role of NEK2 in glioblastoma tumor-initiation, we used 5 lentiviral shRNA clones for NEK2 (shNEK2 D5-D9) and a non-targeting shRNA (shNT: negative control) for g528 and g83 spheres (their sequences are described in Table S1). Western blotting validated the reduction of NEK2 expression at varying degrees with these shRNAs (Fig. S6A). Based on their knock-down (KD) efficiency, we selected shNEK2 D6 and D7 (designated hereafter as shNEK2 #1 (less potent KD) and shNEK2 #2 (more effective KD) (Fig. 4A, Fig. S6A). *In vitro* cell growth kinetics of shNEK2 transduced g528 and g83 glioma spheres was diminished proportionally to the reduction levels of NEK2 by the 2 shRNA clones (Fig. 4B, Fig. S6B). The limiting dilution assay to assess the clonogenic potential of the NEK2 silenced g528 and g83 glioma spheres exhibited substantial decline of the *in vitro* clonogenicity by NEK2 KD (Fig. 4C, Fig. S6C-S6D). The remaining cells in NEK2-silenced g528 glioma spheres appeared to undergo differentiation, since immunocytochemistry of shNEK2-transfected g528 glioma spheres elevated GFAP expression (Fig. 4D).

**Fig. 3.**
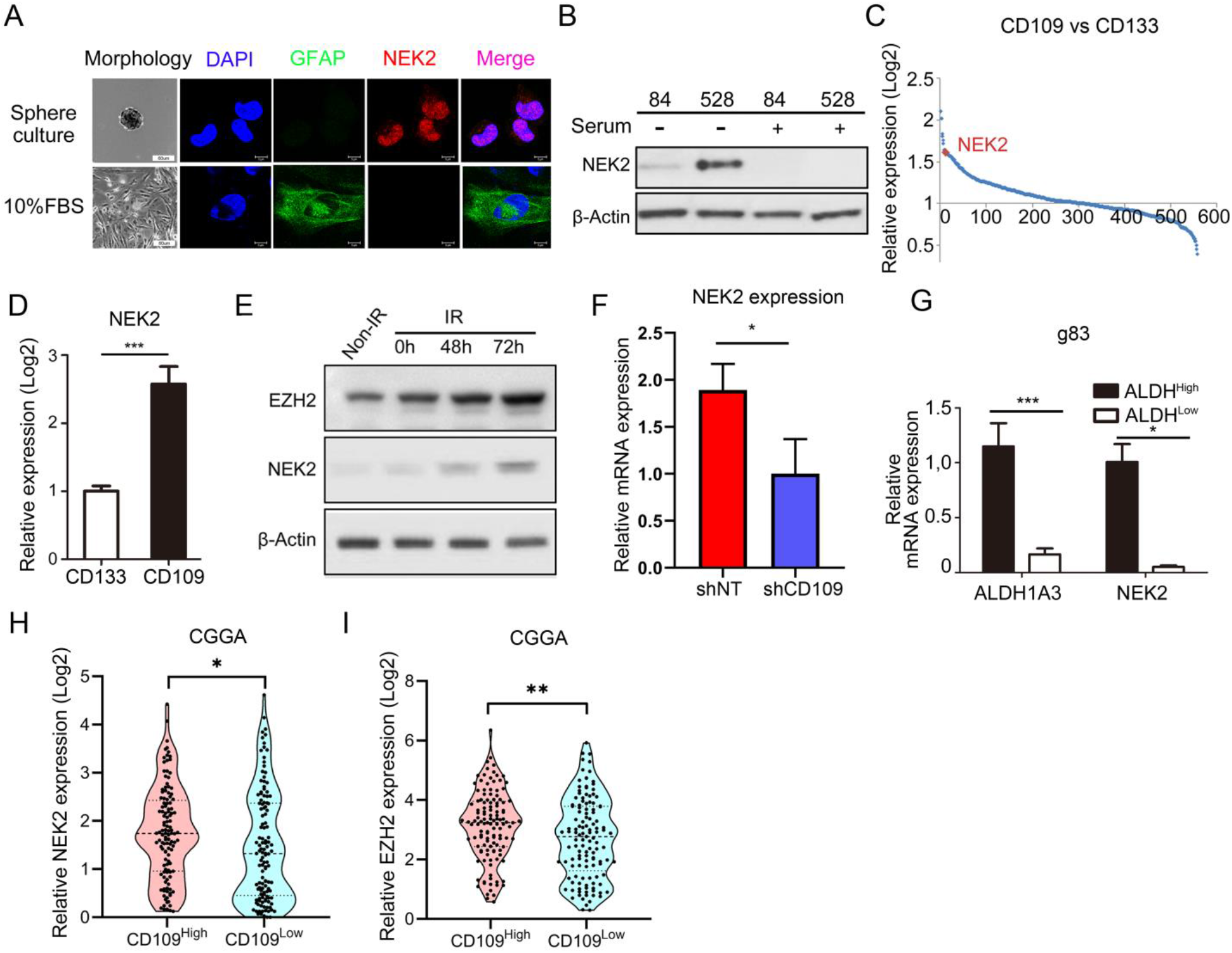
NEK2 expression was associated to CD109-related tumor core signatures and therapy resistance. (A) Immunocytochemistry analysis showed that serum-induced differentiation resulted in reduction of NEK2 expression and elevation of GFAP in g528 spheres. DAPI was used for nuclear staining. (B) Western blotting analysis indicated NEK2 expression was reduced in g528 spheres and g84 spheres exposed to serum for 14 days. β-Actin serves as a loading control. (C) Genome-wide transcriptome microarray analysis showed that *NEK2* was one of the most up-regulated kinase-encoding genes in irradiation-induced CD133-to-CD109 transition glioma sphere. (D) Irradiation (12Gy) was used to induce CD133-to-CD109 transition and qRT-PCR analysis showed that *NEK2* mRNA expression was increased in CD109^High^ g528 spheres compared to CD133^High^ g528 spheres (\*\*\**P* < 0.001, with *t* test). (E) Western blotting analysis showed that NEK2, EZH2 and H3Kme27 expression was increased in g528 spheres treated with irradiation (12Gy) compared with naïve g528 spheres. β-Actin serves as a loading control. (F) RNA sequencing analysis indicated that CD109 knock-down significantly reduced *NEK2* expression in g267 spheres (n = 3, \**P* < 0.05, with *t* test). (G) qRT-PCR analysis showed that *NEK2* mRNA expression was enriched in ALDH^High^ g83 spheres compared to ALDH^Low^ g83 spheres (\*\*\**P* < 0.001, with *t* test). (H) Expression analysis indicated that *NEK2* was significantly enriched in CD109^High^ glioblastoma samples and CD109^Low^ samples in CGGA database (n = 124 for CD109^High^ samples and n = 125 for CD109^Low^ samples, \**P*<0.05, with *t* test). (I) Expression indicated that *EZH2* was significantly enriched in CD109^High^ glioblastoma samples and CD109^Low^ samples in CGGA database (n = 124 for CD109^High^ samples and n = 125 for CD109^Low^ samples, \*\**P*<0.01, with *t* test).

**Fig. 4.**
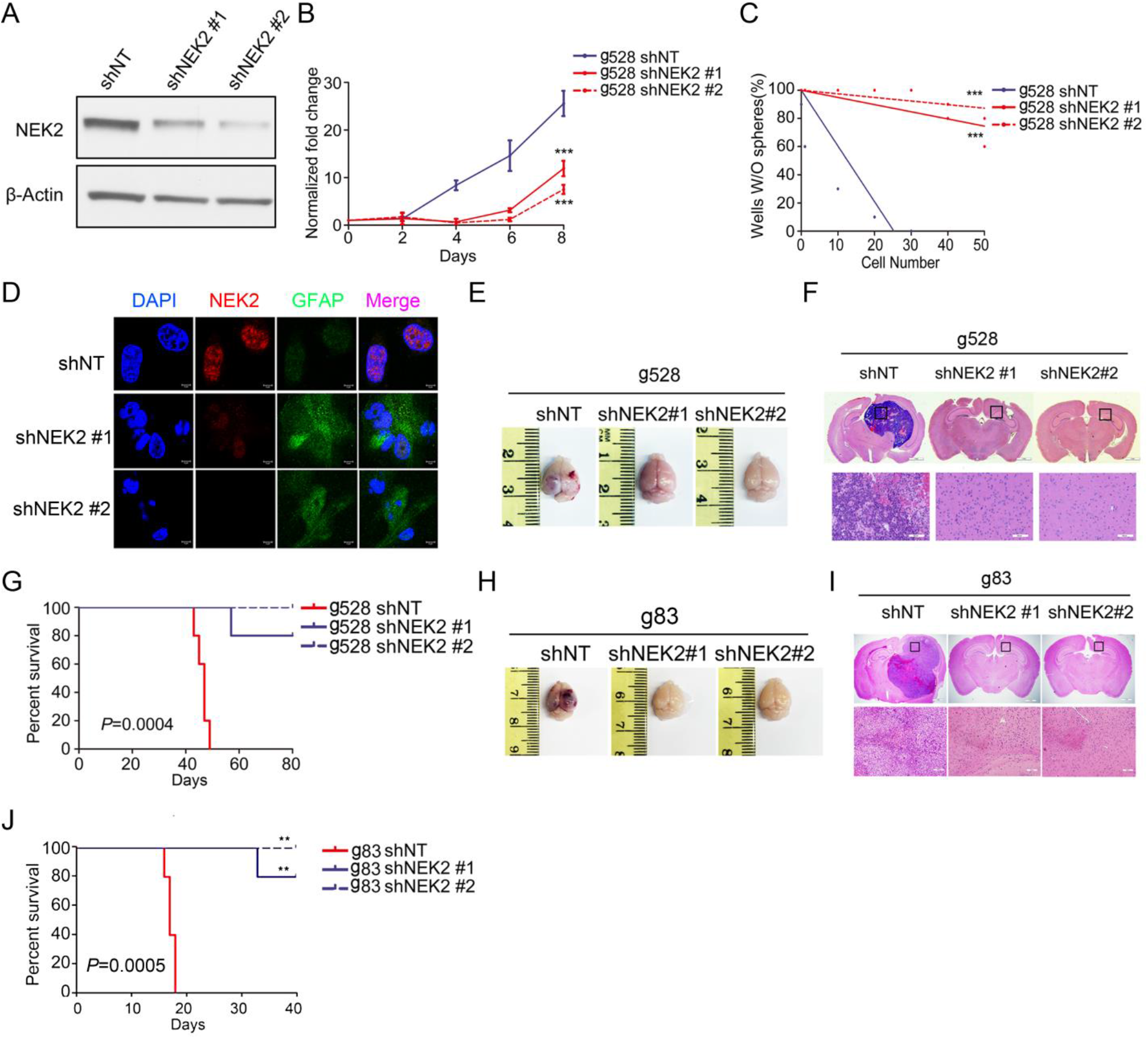
NEK2 silencing reduces tumorigenesis of glioblastoma TICs both *in vitro* and *in vivo*. (A) Western blotting analysis showed that NEK2 expression in g528 spheres were significantly reduced by shRNAs against NEK2 (shNEK2 #1 and shNEK2 #2) compared to non-targeting control (shNT). β-Actin serves as a loading control. (B) *In vitro* growth assay showed that shRNAs against NEK2 (shNEK2 #1 and shNEK2 #2) inhibited cell proliferation of g528 spheres (*P* < 0.0001, n = 6, with one-way ANOVA). (C) *In vitro* clonogenicity assay by limiting dilution neurosphere formation indicated that NEK2 silencing decreased clonogenicity of g528 spheres (*P* < 0.001, with ELDA analysis). (D) Immunocytochemistry analysis indicated that shRNAs-induced silencing of NEK2 resulted in elevation of GFAP expression in g528 spheres. DAPI was used for nuclear staining. (E) Representative images of mouse brains after the intracranial transplantation of g528 spheres transduced with shRNA against NEK2 (shNEK2 #1 and shNEK2 #2) or non-targeting control (shNT). The results showed that NEK2 KD significantly reduced tumorigenesis of g528 spheres *in vivo*. (F) Representative images of H&E stained mouse brain sections after the intracranial transplantation of g528 spheres transduced with shRNAs against NEK2 (shNEK2 #1 and shNEK2 #2) or non-targeting control (shNT). The results showed that NEK2 KD significantly reduced tumorigenesis of g528 spheres *in vivo*. (G) Kaplan-Meier analysis indicated that NEK2 KD markedly prolonged the survival of mice harboring intracranial tumor derived from g528 spheres transduced with shNT (n = 6), shNEK2 #1 (n = 5) and shNEK2 #2 (n = 5). (*P* = 0.0004, with log-rank test). (H) Representative images of mouse brains after the intracranial transplantation of g83 spheres transduced with shRNA against NEK2 (shNEK2 #1 and shNEK2 #2) or non-targeting control (shNT). The results showed that NEK2 KD significantly reduced tumorigenesis of g83 spheres *in vivo*. (I) Representative images of H&E stained mouse brain sections after the intracranial transplantation of g83 spheres transduced with shRNAs against NEK2 (shNEK2 #1 and shNEK2 #2) or non-targeting control (shNT). The results showed that NEK2 KD significantly reduced tumorigenesis of g83 spheres *in vivo*. (J) Kaplan-Meier analysis indicated that NEK2 KD significantly prolonged the survival of mice harboring intracranial tumor derived from g83 spheres transduced with shNT (n = 6), shNEK2 #1 (n = 5) and shNEK2 #2 (n = 5). (*P* = 0.0005, with log-rank test).

Next we investigated the effect of NEK2 KD on *in vivo* tumor formation. Strikingly, while the control mice with xenografts of shNT-transduced g528 or g83 glioma spheres rapidly formed lethal hyper-vascular glioblastoma core-like tumors within 50 days (median survival being 46.2 ± 2.28 days with g528 and 18.2 ± 0.84 days with g83), 4/5 or 5/5 of the mice transplanted with shNEK2 #1- or shNEK2 #2-transduced g528 and g83 glioma spheres failed to form tumors (Fig. 4E-4J), highlighting a potent anti-tumor initiating effect by NEK2 KD. Taken together, NEK2 is essential for survival and growth of the glioblastoma TICs both *in vitro* and *in vivo*.

### NEK2 forms a protein complex with EZH2 to protect EZH2 from proteasome-dependent degradation in glioblastoma

Following these biological data, we sought to understand the molecular mechanism of regulation of EZH2 by NEK2. We first noticed that EZH2 protein expression level is substantially decreased by OTS167 treatment for U87-derived mouse tumors, while the change in *EZH2* mRNA level was rather modest (Fig. 5A-5B). We then performed a protein degradation assay for EZH2 using g267 spheres following 4 weeks of treatment with OTS167 (g267-OTS) at the IC50 dose (10 nM) or vehicle (DMSO) with cycloheximide (CHX) to block protein synthesis. g267-OTS had a longer half-life of EZH2 protein compared to their naïve counterpart (Fig. 5C). We then tested if NEK2 contributes to post-translational regulation of EZH2. With immunoprecipitation (IP) using NEK2 antibody for g528 and g83 spheres, followed by western blotting with EZH2 antibody, NEK2 was found to physically bind to EZH2 in both models (Fig. 5D, Fig. S7A). This physical interaction of EZH2 and NEK2 was confirmed by reciprocal IP by using the same glioma spheres (Fig. 5E, Fig. S7B).

**Fig. 5.**
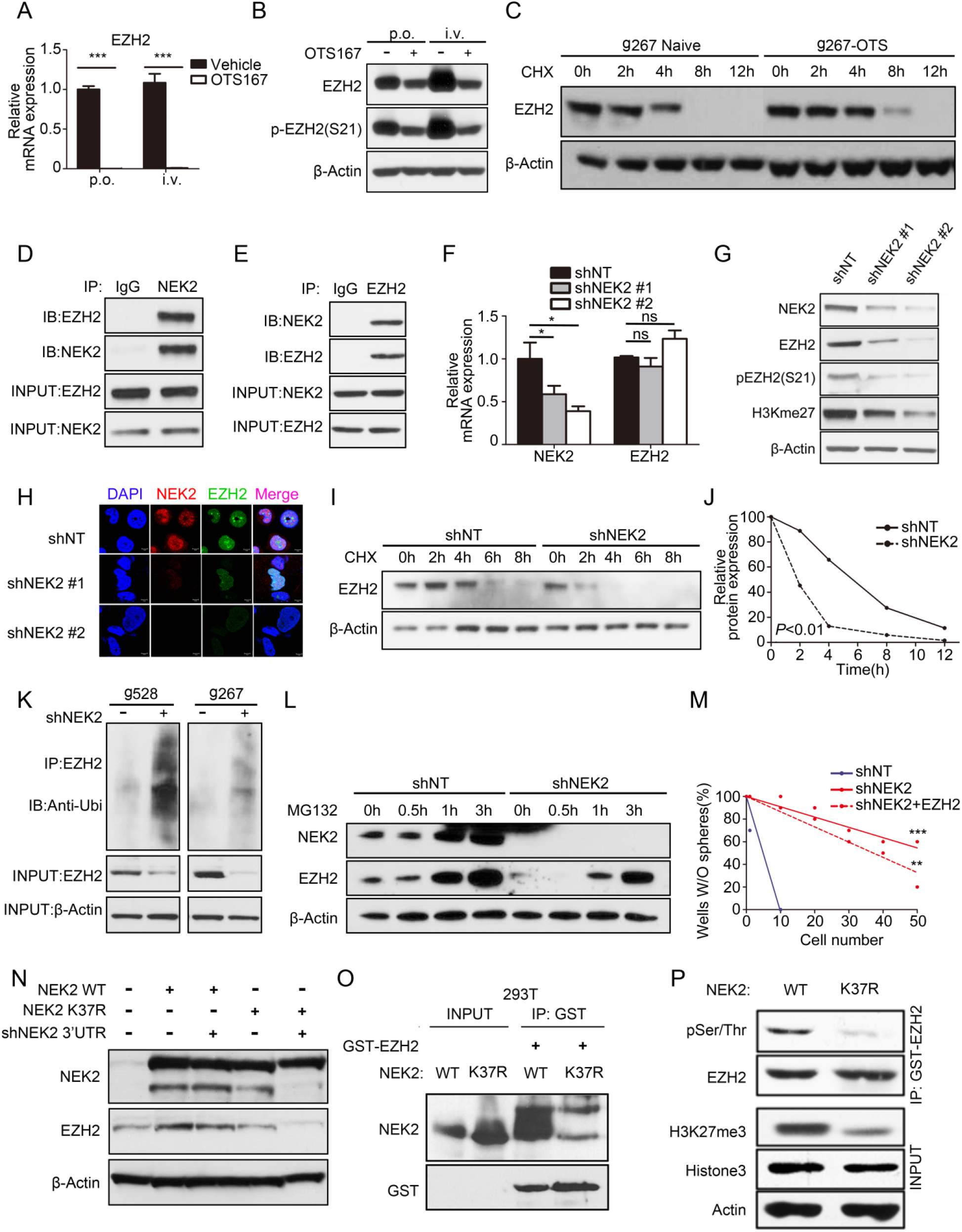
NEK2 signaling is mediated through regulation of EZH2 protein stability in TICs. (A) qRT-PCR analysis showed that EZH2 mRNA expression level was substantially decreased by OTS167 treatment in U87 tumor from sub-Q xenograft mouse model (n = 3, \*\*\**P*<0.001, with *t* test). (B) Western blotting analysis shows EZH2 protein was only modestly affected by OTS167 treatment in U87 tumor from sub-Q xenograft mouse model. β-Actin serves as a loading control. (C) Western blotting analysis for cycloheximide assay of g267 spheres which were pre-treated with OTS167 at IC50 concentration (10 nM) for 4 weeks. The results showed that EZH2 protein stability was enhanced in OTS16t resistant g267 spheres compared to its naïve counterpart. β-Actin serves as a loading control. (D) Western blotting analysis of immunoprecipitation using NEK2 antibody or normal mouse IgG showed that NEK2 physically formed a protein complex with EZH2 in g528 glioma spheres. (E) Western blotting analysis of immunoprecipitation using EZH2 antibody or normal mouse IgG showed that EZH2 physically formed a protein complex with NEK2 in g528 glioma spheres. (F) qRT-PCR analysis indicated that *NEK2* silencing modestly affected EZH2 mRNA expression in g528 spheres (ns *P* > 0.05, \**P* < 0.05, n = 3, with *t* test). (G) Western blotting analysis of NEK2, EZH2, phosphorylated EZH2 at Serine 21(p-EZH2(S21)), trimethyl-HistoneH3 (Lys27) expression were significantly decreased in g528 spheres transduced with shRNAs against NEK2 (shNEK2 #1 and shNEK2 #2). β-Actin serves as a loading control. (H) Immunocytochemistry indicated that EZH2 expression was remarkably reduced in g528 spheres transduced with shRNAs against NEK2 (shNEK2 #1 and shNEK2 #2). DAPI was used for nuclear staining. (I-J) Western blotting (I) and quantization analysis (J) of cycloheximide treated g528 spheres pre-transduced with shRNAs against NEK2 (shNEK2 #1 and shNEK2 #2) or non-targeting control (shNT). The results showed that EZH2 exhibited a shorter half-life in the presence of silenced NEK2. (K) Total EZH2 was immunoprecipitated using EZH2 antibodies from cell lysate of g528 spheres or g267 spheres transduced with shRNAs against NEK2 (shNEK2) or non-targeting control (shNT). The ubiquitinated EZH2 levels were clearly increased in shNEK2 samples compared with shNT samples in both g528 and g83 spheres. (L) Western blotting analysis of EZH2 is performed in MG132 treated g528 spheres transduced with shRNAs against NEK2 (shNEK2 #1 and shNEK2 #2) or non-targeting control (shNT). EZH2 protein underwent degradation after NEK2 silencing and this effect could be partially reversed by proteasome inhibitor MG132. (M) *In vitro* clonogenicity assay by limiting dilution neurosphere formation indicated that NEK2 silencing decreased clonogenicity of g267 spheres and it could be partially rescued by EZH2 overexpression (\*\**P* < 0.01, \*\*\**P* < 0.001, with ELDA analysis). (N) Western blotting analysis indicated that shNEK2 decreased EZH2 expression in g267 spheres and this effect could be rescued by NEK2-WT overexpression vector but not K37R mutation vector. (O) NEK2-WT and mutant (kinase dead mutant: K37R) proteins were subjected to a GST pull-down assay using GST-EZH2 and detected by western blot analysis using anti-NEK2 antibody. The results indicated that EZH2 physically formed a protein complex with wild-type but not K37R mutation type of NEK2 in g022 spheres. Western blotting assay of GST-EZH2 showed that similar amounts of each protein were used. (P) GST-EZH2 was immunoprecipitated using anti GST antibodies from cell lysate of g022 with GST-EZH2 and NEK2-WT or K37R mutation. The results showed that phosphorylation of EZH2 and H3K27me3 was increased by wild-type but not K37R mutation type of NEK2 in g022 spheres.

Next, we investigated whether silencing of NEK2 affects the temporal change of EZH2 expression in glioma spheres. qRT-PCR with shNEK2-transduced g528 spheres showed no noticeable change in *EZH2* mRNA expression with or without NEK2 KD (Fig. 5F). In contrast, western blotting showed that EZH2 protein levels (both the total form and phosphorylated active form) were strongly suppressed by NEK2 silencing in g528 spheres (Fig. 5G, Fig. S7C). The level of H3K27 trimethylation, a downstream target of EZH2, displayed similar results (Fig. 5G). Likewise, immunocytochemistry showed the reduction of EZH2 protein by NEK2 KD in both g528 spheres (Fig. 5H). Furthermore, qRT-PCR detected the downregulation of a set of EZH2 downstream target genes(*25*) by NEK2 silencing in g528 spheres (Fig. S7D). To dissect the molecular mechanisms underlying the NEK2-mediated regulation of EZH2 protein levels in glioma spheres, we evaluated whether NEK2 silencing alters the EZH2 protein expression through inhibiting *de novo* protein synthesis and/or promoting proteasome-mediated protein degradation. When g528 spheres were pre-treated with cycloheximide, EZH2 protein was more rapidly decayed by shNEK2 infection in comparison to shNT control, indicating that NEK2 stabilized EZH2 protein from degradation (Fig. 5I-5J). Consistent with this data, ubiquitinated EZH2 levels were clearly increased in shNEK2 samples compared with shNT samples in both g528 and g83 spheres (Fig. 5K). Western blotting showed that pre-treatment of both g528 and g83 spheres with proteasome inhibitor MG132 exhibited temporal accumulation of NEK2 and EZH2 protein in shNT infected cells, whereas the initial shNEK2-mediated decrease of EZH2 protein was largely, if not completely, reversed by MG132 (Fig. 5L, Fig. S7E-S7H). Collectively, these data suggest that NEK2 is essential to protect EZH2 from proteasome-mediated protein degradation in glioblastoma cells.

We then investigated whether exogenous expression of EZH2 could rescue the effect of NEK2 silencing in g267 glioma spheres. The attenuated proliferation and clonal sphere formation ability resulting from NEK2 KD were both partially rescued by EZH2 overexpression (Fig. 5M, Fig. S7I-S7J). Furthermore, a wound healing assay showed that the ability of cell motility of g267 spheres was decreased by NEK2 silencing and could be rescued by exogenous expression of EZH2 (Fig. S7K). Altogether, function of NEK2 is largely, if not completely, mediated through regulation of the EZH2 signaling in glioblastoma cells.

To clarify whether theNEK2-mediated EZH2 regulation is dependent on its kinase activity, we designed overexpression vectors for NEK2 wild type (WT) and K37R mutation type, which is a known major catalytically inactive mutant of NEK2(*32*). Following transfection of these vectors into g267 glioma spheres, shNEK2 decreased EZH2 expression and this effect was rescued by the NEK2 WT overexpression vector but not by the K37R mutation vector (Fig. 5N). Additionally, NEK2-WT and K37R proteins were subjected to a GST pull-down assay using GST-EZH2 and detected by western blotting using anti-NEK2 antibody in 293T cells and g022 spheres. NEK2-WT could be strongly detected in GST-EZH2 pull down samples, while K37R mutation showed very little association with EZH2 protein (Fig. 5O, Fig. S7L-S7M). Furthermore, when we used three 3’UTR-shNEK2 lentiviral clones (shNEK2 D5, shNEK2 09, shNEK2 48) to block NEK2 expression in g267 spheres, the NEK2 KD-induced decline of proliferation could be rescued by exogenous expression of NEK2-WT overexpression but not K37R mutation type (Fig. S7N-S7P). GST-EZH2 was then immunoprecipitated using anti-GST antibodies from cell lysate of g022 with overexpression of GST-EZH2 and NEK2-WT or K37R mutation. Western blotting exhibited that both EZH2 phosphorylation level and H3K27me3 were dramatically decreased in K37R overexpressing cells compared with NEK2-WT or GST-EZH2 overexpressing cells (Fig. 5P). Altogether, these data indicated that NEK2 regulated EZH2 post-translational expression *via* NEK2 kinase activity.

### Development of a novel NEK2 inhibitor for glioblastoma therapy

In efforts to develop NEK2-targeted therapeutics, we sought to design novel, clinically-applicable small molecules that selectively inhibit NEK2 kinase activity in cancer cells. To this end, we established a NEK2 computational binding model and a focused kinase inhibitor fragment library (~300 compounds) to identify NEK2-selective inhibitors. From the initial screening of the kinase fragment library containing 300 compounds, we identified fragment 1 with activity against NEK2 (IC50 = 20 µM). Using the computer-based drug discovery methodology with fragment 1, we designed and synthesized compound 2, which showed improvement of binding to the NEK2 active site according to computational modeling (Fig. 6A). Compound 2 indeed showed improvement of the NEK2 enzymatic activity with an IC50 of 0.1 μM. Introduction of a methyl group to the benzylic position and replacement of the phenyl linker with a thiophene of compound 2 produced compound 3, which furnished a 3-fold improvement in activity (IC50 = 0.03 μM). Compound 3 was a mixture of two enantiomers, and the pure single isomer 3a from the chiral separation is about 2-fold more potent (IC50 = 0.015 μM) than the racemic mixture 3, while the IC50 of the other isomer 3b is > 0.2 μM. The enantio-pure single isomer 3a or 3b could be obtained from the commercially available R or S 1-(2-(trifluoromethyl)phenyl) ethanol, respectively (Fig. 6A). Therefore, we picked up Compund3a (hereafter designated as CMP3a) as the lead candidate for cancer therapy development (Fig. 6B). The procedures for CMP3a synthesis are described in supplementary materials (Fig. S8A-S8F). To characterize the efficacy of this novel NEK2 inhibitor, we started by investigating the IC50s of CMP3a in different patient-derived glioma sphere models. As expected, sensitivities of glioma sphere cells *in vitro* to CMP3a were correlated with their NEK2 expression levels, while normal human astrocytes were markedly resistant to CMP3a (Fig. 6C). *In vitro* cell free kinase activity assay showed that the NEK2 kinase activity was strongly inhibited by CMP3a at does between 10-100nM (IC50 of 82.74 nM), which was quilt similar to our *in vitro* IC50 doses (Fig. 6D). To assess the specificity, CMP3a was screened at a 15 nM concentration against 97 kinases representing all kinase clusters (KINOMEscan®). There were three kinases (YSK4, FLT3-ITDD835V, FLT3-ITDF691L) featuring > 65% inhibition. Therefore, CMP3a was considered relatively selective to NEK2 inhibition (Fig. 6E, Fig. S9A). CMP3a’s pharmacokinetic (PK) profile was characterized in rats *via* a contract with Pharmacon. With an i.v. dose at 1 mg/kg, CMP3a demonstrated the following PK properties: T1/2 was 1.4 h; Cl was 7304 (ml/ hr/ kg); VOD was 14872 mL/kg; and Auc was 130 hr*/ hr/ kg). Although the AUC was relatively low, most of the drug was likely distributed into tissues based on the high volume of distribution (VOD) (Fig. 6F). We then picked up two HEC1/NEK2 inhibitor (INH1 and INH6) and one PLK4/NEK2 inhibitor (RO3280) to compare the potential efficiency of CMP3a with those on attenuating tumor cell growth (Fig. S9B-S9D). *In vitro* data indicated that CMP3a attenuated glioma sphere cell growth more efficiently compared with other inhibitors in 2 different glioma sphere lines (Fig. S9E-S9F). We also investigated the effect of CMP3a on *in vivo* tumor initiation and propagation. The mouse intracranial tumors derived from two glioma sphere lines (g267 and g374) were treated with CMP3a from day 7 day after xenografting. Treatment was continued for 10 days through tail vein injection at 10 or 20 mg/kg/day. This CMP3a treatment resulted in a decrease of *in vivo* tumor growth and prolonged overall survival of tumor-bearing mice in comparison to the placebo group (Fig. 6G, Fig. S9G-S9H).

**Fig. 6.**
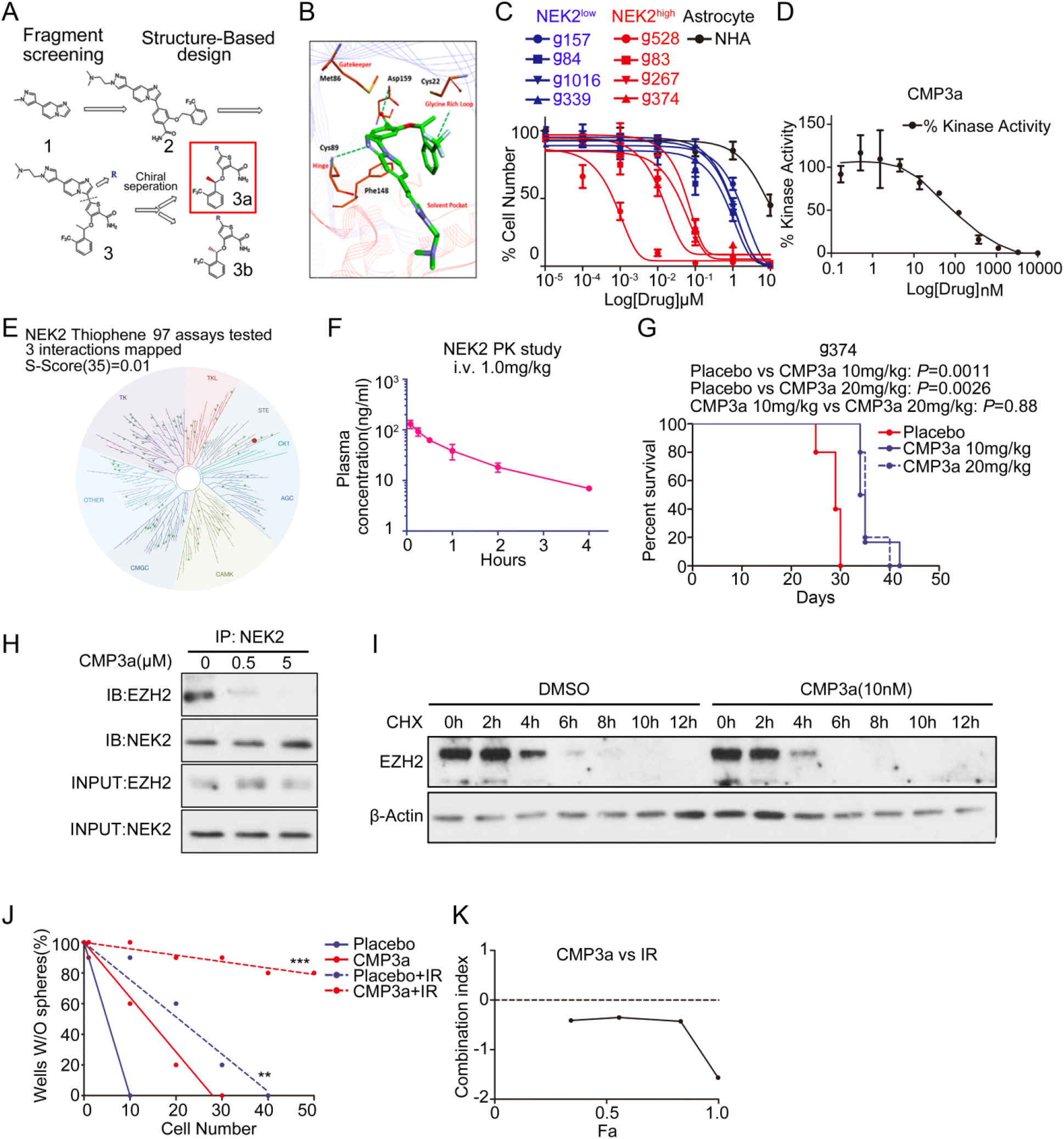
The novel NEK2 inhibitor attenuates tumor growth and increases radio-sensitivity in GBM. (A) The general methods for development of novel NEK2 inhibitor CMP3a. (B) Chemical structure of CMP3a. (C) *In vitro* cell viability assay for CMP3a with NEK2^High^ glioma spheres group (g83, g267, g374 and g528) compared with NEK2^Low^ glioma spheres group (g84, g1016, g339 and g157) and normal human astrocytes (NHA). The results indicated that sensitivity to CMP3a was correlated with NEK2 expression among these glioma spheres. (D) KDs data showed CMP3a blocked NEK2 effectively inhibited NEK2 kinase activity at the dose of 10-100nM. (E) CMP3a was screened at a 15 nM concentration against 97 kinases representing all kinase clusters by using KINOMEscan. The result showed that CMP3a was a relative specific inhibitor of NEK2. (F) Pharmacokinetic data of CMP3a with 1.0mg/ kg intravenous injection in rats. The results indicated that CMP3a was likely distributed into tissues based on the high volume of distribution. (G) Kaplan-Meier analysis indicated that CMP3a treatment prolonged the overall survival of mice transplanted with g374 spheres then followed continuously 10-day CMP3a treatment for different does or placebo by tail vein injection. Data was analyzed with log-rank test. (H) Immunoprecipitation indicated that CMP3a treatment blocked the formation of NEK2-EZH2 protein complex in g528 spheres with or without CMP3a treatment. Western blotting analysis was used to measure EZH2 expression. (I) Western blotting analysis of cycloheximide treated g267 spheres pre-treated with CMP3a or placebo. The results indicated that CMP3a treatment significantly reduced half-life of EZH2 in g267 spheres. (J) *In vitro* clonogenicity assay by limiting dilution neurosphere formation indicated that CMP3a decreased clonogenicity of g528 spheres (*P* < 0.001, with ELDA analysis). (K) CI-Fa analysis showed commination index (CI) of radiation treatment and CMP3a, which indicated that CMP3a had synergistic effects with irradiation on g528 spheres.

Since NEK2 silencing decreased EZH2 expression by destabilizing EZH2 protein, we treated glioma spheres with CMP3a for 48h with varying doses. Western blotting indicated that both EZH2 and H3K27me3 expression were reduced in a dose-dependent manner with CMP3a (Fig. S9I). We then purified protein from g528 spheres treated with or without CMP3a (0.5μM and 5μM respectively) for 2h, followed by immunoprecipitation with NEK2 antibody. Western blotting showed that treatment with CMP3a resulted in a blockage of the NEK2-EZH2 complex (Fig. 6H). Additionally, a cycloheximide assay yielded that the half-life of EZH2 protein was decreased in g528 spheres after 24h treatment with CMP3a, which represented a consistent result with shNEK2 KD (Fig. 6I).

As NEK2 has been shown to be a mitotic kinase, promoting tumor cell proliferation mainly through centrosome separation and bipolar spindle formation at centrosomes(*33*), we sought to clarify whether the function of NEK2 is mainly the regulation of glioblastoma cell mitotic progression. To test this, immunocytochemistry was performed and acetylated α-Tublin was used to label the micro tubes and centrosomes. With confocal microscope, we observed that NEK2 protein was still localized to centrosomes after treatment of CMP3a (Fig. S9J) and the mitosis of glioma sphere cells was not largely affected by CMP3a treatment (Fig. S9K). Additionally, cell cycle analysis exhibited that the expression levels of NEK2 in glioma spheres were comparable in all phases of cell cycle including the mitotic phase (Fig. S9L). Flow cytometry also showed that inhibition NEK2 by CMP3a did not exhibit any noticeable cell cycle changes of glioma spheres at IC50 dose (Fig. S9M).

To move one step further toward clinical application, we combined CMP3a treatment with radiation using the g528 sphere model. Western blotting indicated that EZH2 was increased after radiation, which was eliminated by NEK2 inhibition by using CMP3a (Fig. S10A). When we combined CMP3a treatment with radiation treatment in g528 glioma spheres, CMP3a treatment decreased radio-resistance in g528 glioma spheres (Fig. 6J, Fig. S10B-S10C). Furthermore, flow cytometry showed radiation treatment increased the proportion of apoptotic cells in a dose-dependent manner, when combined with CMP3a (Fig. S10D-S10E). To distinguish whether NEK2 inhibition had an additive or synergistic effect with radiation, the Chou-Talalay model was applied to the obtained data. The data showed combination index (CI) of irradiation and CMP3a was less than 1, which means NEK2 inhibition has synergistic effects with radiation on g528 spheres (Fig. 6K).

Additionally, we used NHA to get an idea for the safety of CMP3a combined with radiation. Unlike glioblastoma cells, qRT-PCR data showed that NEK2 mRNA expression was dramatically decreased after radiation (Fig. S10F) and CMP3a did not significantly affect sensitivity of NHA cells to radiation (Fig. S10G). Altogether, these data suggested that the NEK2 inhibitor is capable of decreasing glioblastoma growth and its radio-resistance through regulating EZH2 protein stability.

## Discussion

Successful establishment of the tumor core results in a harsh microenvironment, necessitating a subset of glioblastoma cells to undergo molecularly dynamic changes(*2*). Similarly, successful tumor recurrence following failure of multi-modal therapies invokes cancer cell response, although these spatial and temporal changes in tumors may not be exactly the same(*3*). Therapy-resistant tumor core cells had high EZH2 expression, and inhibitors were more effective in attenuating growth of the tumor core-derived models both *in vitro* and *in vivo*. Yet, the differences were relatively modest. On the other hand, increased elevation of NEK2 within the tumor core cells was considerably pronounced. Longitudinally, NEK2 activation emerged through long-term treatment with OTS167 to sustain EZH2 function, elevating therapy resistance in glioblastoma. Given the elevated expression of NEK2 in the tumor core as opposed to the edge, and in recurrence contrary to primary tumors, NEK2-driven EZH2 persistence may be a mechanism that enables glioblastoma tumors to spatio-temporally adapt to challenging conditions (Fig. 7). However, several questions remain. First, the molecular signal that triggers this evolving NEK2 activation is undetermined. In addition, the role of NEK2 in tumor initiation was determined from tumor core models. It remains unclear whether this means NEK2 regulates the continuous E-to-C progression, or solely maintains the expansion of established tumor core cells. Future experimentation will address these remaining questions. Furthermore, the extent of possible resection of tumor edge and core varies between individual tumors depending on their proximity/invasiveness to neighboring functional areas. Therefore, core-targeting (or E-to-C targeting) therapies should be combined with edge-targeting approaches to achieve clinically meaningful benefit for glioblastoma patients.

**Fig. 7.**
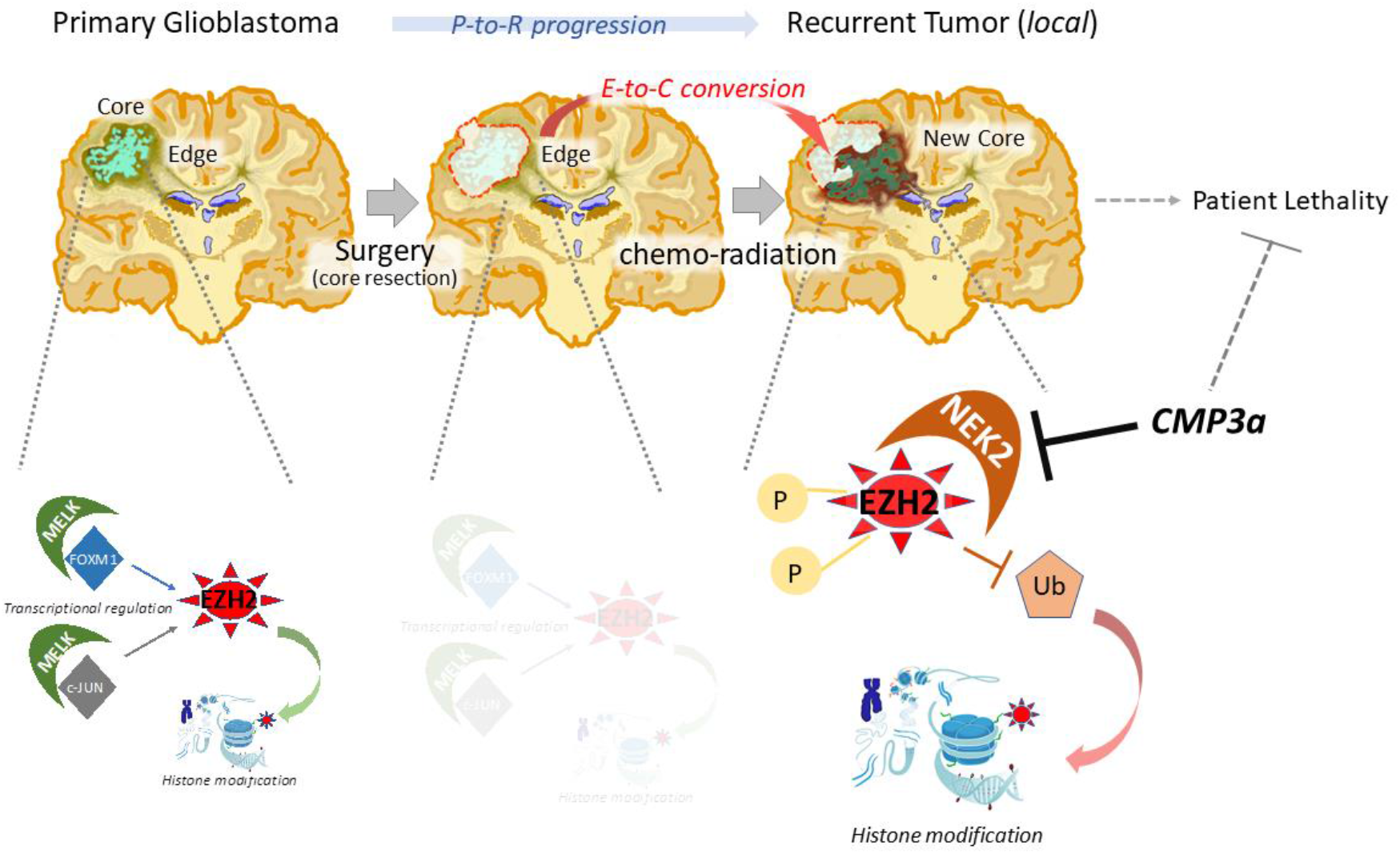
NEK2-dependent post-translational regulation of EZH2 is essential for edge-core conversion thus induced therapy resistance in glioblastoma.

Protein kinases represent a druggable molecular target in a number of diseases and play a fundamental role in various therapy-refractory cancers(*34*). However, rapid and dynamic resistance to “key” kinase inhibitions is frequently observed, leading to subsequent treatment failure(*34*). From a therapeutic standpoint, our target disease state is, in fact, the lethal recurrence following therapeutic failure, rather than the treatment-naive primary tumor(*35*). Thus, identifying the dynamics in glioblastoma cells following therapeutic insult needs deeper investigation. NEK2 is a poorly-characterized gene, encoding a serine/threonine kinase(*32*). Its elevated expression has been reported in various cancers including breast, lung, colorectal cancers and glioma(*36–40*). Higher expression of NEK2 is correlated with poorer prognosis of glioma patients(*40*). Nonetheless, no clinical trials for NEK2 inhibition have been designed in any cancers. Current attempts at developing a NEK2-targeting strategy with CMP3a provided encouraging results in pre-clinical models. Caution is paramount, however, as CMP3a or its derivatives may induce further escape of some glioblastoma cells that retain (or provoke) recurrence-initiating potential. The synergistic effect of CMP3a with radiation may guide us toward clinical trial development, although the molecular mechanism for this synergism needs further investigation. For successful therapeutic development for glioblastoma with CMP3a and/or other targeting agents, it is crucial to further understand the temporal and spatial changes within the heterogeneous cancer cell populations that allow cancers to evade any given treatment.

## Acknowledgments

We would like to express our sincere appreciation to all the patients and families, who kindly allowed us to obtain tumor samples for this study. To achieve the best benefit for neuro-oncology research on behalf of our patients, we would be more than willing to share our patient-derived experimental models described in this study with any scientists who express interest, once the appropriate paperwork is agreed. We would also like to thank all of our collaborating scientists, as well as the assigned reviewers, editors, and staff members in the editorial office for their constructive comments and administrative help in this manuscript. A part of this work was previously published in the Journal of Clinical Investigation (Wang et al., 2017), yet due to some questionable data, we retracted the previous paper, repeated all experiments that presented any concern with extensive additional experimentation to obtain further confirming data, and created a new manuscript (see more information in supplementary information including Fig. S11 and Fig. S12). We acknowledge the contribution by all the members in the Nakano laboratory (past and present) for technical help. This work was supported by the NIH grants R01NS083767, R01NS087913, R01CA183991, and R01CA201402 to IN.

## Author contributions

Leading conceptualization of the study: IN. Financial support: IN, HYL. Overall design of the study: JW, MSP, IN, with input from DY, YS, YL. Laboratory practice: JW, MSP. Analysis of data; JW, MSP, IN. H.L. Design, synthesis, and screening of NEK2 inhibitors, *in vitro* KINOMEscan^®^ and PK analysis: BF, WHH, YHS, HYL. shRNAs for NEK2: VG. Drafting the article: JW, IN. Critical revision of the article: MAN, IN. All authors had substantial input to the logistics of the work and revised and approved the final manuscript. The authors know their accountability for all aspects of the study ensuring that questions regarding the accuracy and integrity of any part are appropriately investigated and resolved. The corresponding author had full access to all of the data and the final responsibility to submit the publication.

## Materials & Methods

### Patients, Specimens, and Ethics

This study was started and completed under the approved Institutional Review Board (IRB) and Institutional Animal Care and Use Committee (IACUC) protocols in University of California Los Angeles (UCLA), MD Anderson Cancer Center (MDA), University of Alabama at Birmingham (UAB), and Ohio State University (OSU). The IRB Protocol number at UAB is #N151013001.

For the pre-clinical studies, 21 patient-derived glioma sphere models were used, including 3 pair of tumor core- and edge-derived ones (g0573, g1053, g1051), which were established and described elsewhere. In short, the senior author (IN) performed supra-total resection of glioblastoma tumors under the awake setting and resected both tumor core (T1-Gadolinium(+) tumors) and edge (T1-Gadolinium(-)/T2-FLAIR abnormal tumors in the non-eloquent deep white matter) to achieve maximal tumor cell eradication without causing any permanent major deficit in the patients (Fig. 1E). After the confirmation of enough tumor tissues from both lesions secured for the clinical diagnosis, remaining tissues were provided to the corresponding scientists following de-identification of the patient information. Both the core-derived and edge-derived glioma spheres were established in the same culture condition and their spatial identities, termed core-ness and edge-ness, were confirmed by a set of xenografting experiments into mouse brains (details described in Fig. S2B). Only those that passed this confirmation were used for this study. Other glioma specimens and normal brain tissue samples were collected in the Department of Pathology and Laboratory Medicine at UCLA (sample g157, g374, g339), MDA (sample g267, g28, g30, g11, g711), and OSU (sample g83, g528, g84, g57, g185A, g1016 and g2313), and processed to the research laboratories after de-identification of the samples, as described previously(*16, 18–20*). All these patient-derived glioma models were periodically checked with the mycoplasma test and the Short Tandem Repeat (STR) analysis.

### Data reliability and validity

All the experiments were performed by two or more researchers involved in each procedure. The *in vitro* experiments were repeated at least three independent procedures to obtain valid data. All the raw data are placed in Fig. S13 and Fig. S14. The corresponding author (IN) confirmed the agreement of everything related to this study with all the involved authors.

### Statistical analysis

All the data were presented as mean ± SD. The number of replicates for each experiment was stated in the figure legend. Statistical differences between two groups were evaluated by two-tailed *t* test. The comparison among multiple groups were performed by one-way analysis of variance (ANOVA) followed by Dunnett’s posttest. The statistical significance of Kaplan–Meier survival plot was determined by log-rank analysis. Statistical analysis was performed by Prism 6 (Graphpad prism), unless mentioned otherwise in figure legend. *P* < 0.05 was considered as statistically significant.

Other detailed materials and methods can be found in supplementary materials and methods.

