## Supplemental Information for "Spatiotemporal Dynamics of Intra-tumoral Dependence on NEK2-EZH2 Signaling in Glioblastoma Cancer Progression"

***Supplementary Information***

The original version of this study was partially published in the Journal of Clinical Investigation (JCI) in 2017 (2017;127(8):3075–3089.). However, in 2020, the Journal Office notified to the corresponding authors with 5 pieces of the presented data that raised concern. After the extensive investigation within the Nakano laboratory, we found that, although in the first submitted manuscript to JCI contained all the correct data (see Fig. S11), the errors were introduced during the preparation of the revision. The corresponding author (IN) of this manuscript confirmed and clearly affirms here that none the current authors in this manuscript were involved in the conduct of the data replacement that happened in the previous article. Importantly, the fact that the correct data were included in the first submission was confirmed by the Journal Office for JCI, as well. Following extensive discussion with the JCI Journal Office, we decided to retract the previous article from JCI to ensure the validity and reproducibility of this study. After re-confirmation, all five pieces of the correct data are included in this manuscript (Fig. 2D, Fig. 3E, Fig. 4A, Fig. 5G, Fig. S7C). We also extensively performed additional experiments to obtain a set of updated data to improve the quality and logistics of this study. The correct data was shown in Fig. S11 together with the replications and the incorrect bands from the original publication. All the related uncropped gels could be found in Fig. S12.

***Supplementary Materials and Methods***

***Reagents and Antibodies***

Following reagents and primary antibodies are used in this study: EGF (Peprotech), bFGF (Peprotech), B27 (invitrogen), Heprin (Sigma), DMEM-F12 (Gibco, 10565-018), Fetal bovine serum (Gibco, 10082-147), Albumin from bovine serum (Sigma, A2153), Accutase solution (Sigma, A6964-100), alamar Blue (Invitrogen, DAL1100), RIPA buffer (Sigma, R0278), Phosphatase inhibitor cocktail (Sigma, P0044), Protease inhibitor cocktail (P8340), Bradford (BIORAD, 500-0006), BSA used in Bradford assay (BioLabs, B9001S), PageRuler plus prestained protein (Thermo scientific, 26619), iScript Reverse Transcription supermix for RT-qPCR (Bio-rad, 170-8841), Protein A/G Magnetic Beads (Thermo Scientific Pierce, 88802), IP Lysis Buffer(Thermo Scientific Pierce, 87787). anti-NEK2(Abcam,ab117553, Mouse, IHC and IP), anti-NEK2(Thermo Scientific Pierce,PA5-31259, Rabbit, ICC, WB and IF), anti-NEK2(Thermo Scientific Pierce,PA5-15337, Rabbit, WB), anti-EZH2(Cell Signaling, #2146, Rabbit, IP and WB), anti-EZH2(Thermo, MA5-15101, Mouse, ICC and IF), anti-H3K27me3(EMD Millipore, #07-449, Rabbit), β-Actin(Sigma, A 5316, Mouse), anti-Ubiquintin(Cell signaling, #3936, Mouse), anti-Acetylated alpha Tubulin (Abcam, ab24610, Mouse). anti-Ki-67 antibody (Thermo Fisher, 50-5699-82, Mouse), anti-CD45 antibody (LifeSpan BioSciences, LS-C140427, Mouse).

***Drug Treatment and cell viability assay***

Tazemostat and CPI-1205 were obtained from Selleckchem (Houston, TX). Stock solutions for both compounds were prepared using dimethyl sulfoxide (DMSO)

(Sigma Aldrich, St. Louis, MO). In each experiment, DMSO alone was used as a control at a concentration of between 0.1% and 1% (i.e. identical to the DMSO concentrations in drug-treated cells); the growth of the cells used in this study was not significantly affected by DMSO at the concentrations employed. Viability of glioblastoma cells was determined using an AlamarBlue assay (Thermo Fisher Scientific, Rockford, IL). Briefly, cells were seeded at 3,000 cells per well in 96 well plates (excitation at 515-565nm, emission at 570-610nm); a Synergy HTX multi-mode reader (BioTek; Winooski, VT) was utilized for all experiments.

### *Differential gene expression analysis*

Transcriptome profile of GSE 68029 and GSE 56937 were extracted from Gene expression omnibus (GEO) then preprocessed. Gene expression datasets of TCGA GBM was extracted from GDC Data Portal (https://portal.gdc.cancer.gov/). The limma package was used for identifying the differentially expressed genes in these datasets(*1*). The expression difference of individual gene was defined by $Log_2$(Fold change) and adjusted *P* value, in which $log_2FC > 2.0$ with an adjusted $P$ value $< 0.05$ was defined as an up-regulated gene. Venn diagram was carried out to illustrate the overlapping down-regulated genes in these datasets.

### *In vitro cultures*

Core and edge tissues of glioblastoma were identified and resected by intra-operative MRI. Glioblastoma tissues were disassociated into single cells immediately after surgical resection with Accutase at 37°C then transferred into DMEM/F12 medium (Invitrogen) containing 2% B27 supplement (Invitrogen) (vol%), 2.5 mg/ml heparin, 20 ng/ml basic fibroblast growth factor (bFGF, Peprotech), and 20 ng/ml epidermal growth factor (EGF, Peprotech). bFGF and EGF were added twice a week and the culture medium was replaced every 10 days. Experiments with neurospheres were performed with lines that were cultured less in than 40 passages since they initially established. The human fetal neural stem cell sample (16wf) was established at UCLA as described previously(*2, 3*). Normal Human Astrocytes (NHA, Lonza) were used as a control sample in this study.

***Western blotting***

The cell lysates were prepared in RIPA buffer containing 1% protease and 1% phosphatase inhibitor cocktail (Sigma Aldrich) on ice. The sample protein concentrations were determined by the Bradford method. Equal amounts of protein lysates (10 μg/lane) were fractionated on NuPAGE Novex 4-12% Bis-Tris Protein gel (Invitrogen) and transferred to a PVDF membrane (Invitrogen). Subsequently, the membranes were blocked with 5% skimmed milk for 1 h and then treated with the relevant antibody at 4 ℃ overnight. Protein expression was visualized with Amersham ECL Western Blot System (GE Healthcare Life Sciences). β-Actin served as a loading control. Image J was used to analyze the Western blotting results.

***In vivo intracranial xenograft tumor models***

Six-week-old nude mice were used for glioma sphere intracranial xenotransplantation. All animal experiments were carried out under an Institutional Animal Care and Use Committee (IACUC)-proved protocol according to NIH guidelines. The glioblastoma cells suspension (5000 cells for g83 spheres or $1\times10^5$ for g528 spheres or $1\times10^5$ for g267 spheres in 2 μl of PBS) transduced with non-target or shNEK2 lentivirus was injected into the brains of nude mice as previously described(*4*). Six mice were used for each group. When neuropathological symptoms developed, mice were sacrificed and followed by perfusion with ice-cold PBS and 4% (wt/ vol) paraformaldehyde (PFA). Then the mice brains were dissected and fixed in 4% PFA for 24 h and then transferred to 10% formalin and then performed to make sections. Drug treatment was done through tail vein injection.

***Immunoprecipitation***

Lysis the cells with 500μL Lysis Buffer containing protease inhibitor and phosphatase inhibitor. Pick 25ul magnetic beads and wash with 175μL TBST. Then mix all the cell lysis with pre-washed magnetic beads and adjust the volume to 500μL with lysis buffer and incubate with mixing under 4℃ for 1 h to remove unspecific bindings. Collect the supernatant with magnetic stand then incubate with the antibody and mixing under 4℃ overnight. Then move the antigen sample/antibody mixture to the tube and incubate with mixing under room temperature for 1 h. Collect the magnetic beads with magnetic stand and incubate with low-pH Elution at room temperature with mixing for 10 min. Magnetically separate the beads and save the supernatant containing target protein. Normalize the pH with 15μL of Neutralization Buffer for each 100 ul of eluate. Immunoprecipitation samples were running electrophoresis with 4-12% Bis-Tris Protein gel. Sliver staining was performed with Pierce Silver Stain Mass Spectrometry Kit (Thermo, 24600).

***Protein Expression, purification and GST pull down assay***

GST-fusion constructs of EZH2 or GST proteins were transformed into E. coli BL21 (DE3) pLysS cells (Invitrogen, Carlsbad, CA, USA) and grown to an A600 of 0.4. Following the induction with 0.1 mM isopropyl-β-D-thiogalactopyranoside (IPTG) for 6 h at 32 ℃, the fusion proteins were purified using the standard methods (5). Also GST-EZH2 protein was from SignalChem (E396-30G-20). The bacterias were harvested and resuspended in STE buffer (150 mM NaCl, 10 mM Tris–HCl, pH 8.0, 1 mM EDTA, 0.1 mg/ml lysozyme), and incubated on ice for 15 min. Sonication was followed by addition of 100 μl of 1 M DTT and 1.4 ml of 10 % Sarkosyl to 10 ml of cell suspension. The cell lysate was then incubated with 4 ml of 10 % Triton X-100 for 30 min at room temperature, and glutathione-sepharose beads were added and incubated for 4 h at 4 ℃. A complex of the GST-fusion proteins and the beads were obtained by centrifugation.

For GST pull-down assay, purified NEK2-WT and mutant (kinase dead mutant: K37R) were incubated with GST fusion protein-coupled glutathione beads for 4 h at 4 ℃. The precipitates collected by centrifugation were resuspended in 2× Laemmli sample buffer and subjected to SDS-PAGE and Western blot analysis with either anti-NEK2 antibodies.

***Lentivirus Production and Transduction***

Plasmid DNA preparations were obtained using a large-scale plasmid purification kit (QIAGEN and Roche). HEK293FT packaging cells were co-transfected with the pLKO.1 vector encoding the shRNA and the helper plasmids for virus production (psPAX2 and pMGD2), using Trans-IT (Mirus). Before transduction, spheres were dissociated into single cells with accutase. Cells were seeded in Laminin pre-treated 6 cm petri dishes with $4\times10^5$ cells per well in a final volume of 5 ml. Lentivirus contained medium was collected at time points 48h and 72h then add Lenti-Concentrator and incubated with gently mixing at 4 ℃ for 72 h. Produced lentiviruses were concentrated by ultracentrifugation of the HEK293T supernatant at 25,000 rpm. To assess transduction efficiency, cells on each plate were infected with lentivirus expressing green fluorescent protein (GFP) as a control. Transduction was deemed efficient if > 70% cells were GFP-expressing. The sequences of all the plasmid was showed in Table S1.

***RNA Isolation and Quantitative Real-Time PCR***

RNA was isolated by using RNeasy mini kit (QIAGEN) according to the manufacturer’s instructions. RNA concentration was determined using a Nanodrop 2000 (Thermo scientific). cDNA was synthesized by using iScript reverse transcription supermix for qRT-PCR (Bio-rad) according to the manufacturer's protocol. The reverse-transcribed cDNA was analyzed by quantitative RT-PCR (qRT-PCR), and GAPDH or 18s was used as an internal control. Each qRT-PCR included a 10μL reaction mixture per well that includes 2.5 μL cDNA, 0.5 μL forward primer (0.5 μM), 0.5 μL reverse primer (0.5 μM), 1.5 μL of DNase/ RNase-free distilled water, and 5 μL SYBR green reagent (QIAGEN). The following cycles were performed during DNA amplification: 94 ℃ for 2 min, 40 cycles of 94 ℃ (30 s), 60 ℃ (30 s), and 72 ℃ (40 s). All the primer sequences were showed in Table S2.

***Immunohistochemistry staining***

IHC were performed as previously described(*4*). For IHC, experimental mice were sacrificed and perfused with ice-cold PBS followed by 4% (wt/vol) paraformaldehyde (PFA). Then brains were harvested and fixed in 4% (wt/vol) PFA for 24 h and then transferred into 10% formalin. After tissue embedding and sectioning were performed, brain sections were incubated with the indicated primary antibodies overnight at 4 ℃, followed by incubation with an HRP-conjugated secondary antibody for 1 h at room temperature. Signals were detected using DAB substrate kit (Vector). Nuclei were counter stained with hematoxylin or Hoechst, respectively. Samples incubated without primary antibodies were used as negative controls.

***Immunocytochemistry***

ICC methods were described in previous papers(*4*). For immunocytochemistry, neurospheres were dissociated into single cells and seeded onto coverslips coated with 0.5% laminin (2×104 per well). After 24 h, the cells were fixed with 4% (wt/vol) PFA, blocked with 1% BSA containing 0.3% Triton-X, and treated with primary antibody at 4℃ overnight. The cells were then incubated with Alexa Flour 488-conjugated secondary antibody for 1 h at room temperature. The coverslips were mounted with mounting medium containing DAPI (Vecter, H1200). Images were captured using a fluorescence upright microscope (Olympus DP71). Identical filters, objectives, and acquisition parameters were used for each experiment.

***Immunofluorescence and imaging***

Immunofluorescence was done using the protocol described previously(*4*). The immunostained samples were imaged using the Olympus Fluoview 1000 confocal microscope and edited using Photoshop CS6. Clone area and brain lobe size were measured (in pixels) for quantification by using the Histogram function of Adobe Photoshop CS6.0. Data presented shows mean values and error bars based on standard deviation calculated for each sample.

***PCR-mediated Mutagenesis***

The PCR-mediated mutagenesis method was used to obtain the NEK2 mutant K37R. The primer pairs P1/P4 and P3/P2 were used to amplify two NEK2 fragments. The PCR mixture was incubated first at 95℃ for 5 min, followed by 95℃ denaturing for

30 second, 60 ℃ annealing for 30 second, and 72 ℃ extension for 1 min. After 18 cycles, the PCR1 products were mixed and used as template in a second PCR (PCR2), primed by oligonucleotides P1 and P2 to carry out another 18 cycles of PCR with the same PCR conditions used in PCR1. Resulting PCR2 product was digested with BstBI/BamHI and cloned into corresponding sites of pCDH-EF1-MCS-IRES-Puro plasmid (System Biosciences). Obtained plasmid was designated as pCDH-mutNEK2. The absence of unwanted mutations in the inserts and vector-insert boundaries was verified by sequencing. The sequences of 3'UTR-shRNAs and primers used in this experiment are provided in Table S1 and Table S3.

***General Compound Synthetic Experiments***

All solvents were reagent grade or HPLC grade and all starting materials were obtained from commercial sources and used without further purification. Purity of final compounds was assessed using a Shimadzu ultra-high throughput LC/MS system (SIL-20A, LC-20AD, LC-MS 2020, Phenomenex® Onyx Monolithic C-18 Column) at variable wavelengths of 254 nM and 214 nM (Shimadzu PDA Detector, SPD-MN20A) and was > 95%, unless otherwise noted. The HPLC mobile phase consisted of a water-acetonitrile gradient buffered with 0.1% formic acid. 1H NMR spectra were recorded at 400 MHz and 13C spectra were recorded at 100 MHz, both completed on a Varian 400 MHz instrument (Model# 4001S41ASP). Compound activity was determined with the EZ Reader II plate reader (PerkinElmer®, Walthman, USA). All compounds were purified using silicagel (0.035-0.070 mm, 60 Å) flash chromatography, unless otherwise noted. Microwave assisted reactions were completed in sealed vessels using a Biotage Initiator microwave synthesizer.

***Synthesis of Compound 2 (C2)***

4-chloropyridin-2-amine (7.425 g, 57.8 mmol) and 2-chloroacetaldehyde 50% w/w (8.80 mL, 69.3 mmol) were heated in 1-butanol (23.10 mL, 2.5 M) at 130 ℃ for 12 hours. The solvent was evaporated from the reaction and the title compound was purified *via* DCM/MeOH and isolated as an off-white solid, 1a (6.95 g, 79%). 1a (35.6 g, 233 mmol) was dissolved in acetic acid (292 mL). The reaction was stirred under ice and bromine (13.85 mL, 268 mmol) was added dropwise. The resulting precipitate was filtered off and neutralized to generate the free-base, brominated compound, 1b (35 g, 64.8%). 4-chloro-2-hydroxybenzoic acid (30 g, 174 mmol) was added to methanol (60 mL) and sulfuric acid (130 mmol, 6.94 mL). The reaction was stirred at reflux for 120 hours. The reaction was neutralized with the addition of solid NaHCO3, diluted with water, and extracted with 6:1 ether/DCM. The organic was dried with MgSO4 and condensed to generate the title compound, 1c (31.4 g, 97%). 1-(2-(trifluoromethyl) phenyl) ethanone (19.92 mL, 133 mmol) was dissolved in THF (400 mL). Following, sodium borohydride was (7.37 g, 199 mmol) was added and the reaction was heated to 55 ℃ for 48 hours. The reaction was quenched with the addition of dilute HCl under ice. The organic was extracted with ethyl acetate, dried with MgSO4, and isolated as a viscous oil, 1d (24 g, 95%). 1c (5 g, 26.8 mmol) and 1d (5.61 g, 29.5 mmol) were dissolved in DCM (80 mL) and cooled in an ice bath.

Following, triphenylphosphine (10.18 g, 38.9 mmol) was added and DEAD (6.97 g, 38.9 mmol) was added dropwise. After the addition, the reaction was removed from the ice bath and stirred at room temperature for 12 hours. The crude reaction was condensed and adsorbed onto silica and purified *via* flash chromatography using Hexanes/EtOAC. The title compound was isolated as white solid/crystals, 1e (6.12 g, 63.7%). 1e (6.122 g, 17.07 mmol) was dissolved in dioxane (100 mL) and degassed with argon. Following 4,4,4',4',5,5,5',5'-octamethyl-2,2'-bi(1,3,2-dioxaborolane) (6.50 g, 25.6 mmol), potassium acetate (5.02 g, 51.2 mmol), Pd2(dba)3 (0.156 g, 0.171 mmol), and PCy3 (0.143 g, 0.512 mmol) was added to the reaction. The reaction was placed under an argon atmosphere and heated to 100 ℃ for 12 hours. The crude reaction was added to water, extracted with ether, and washed with water 5x. The ether extract was collected, dried, and absorbed onto silica. The reaction was purified *via* flash chromatography using hexanes/EtOAc to generate the title compound as a yellow oil, 1f (7.47 g, 97%). 1b (0.615 g, 2.66 mmol) and 1f (1.196 g, 2.66 mmol) were added to 4:1 DMF/Water (26 mL). The reaction was degassed with argon and bis(triphenylphosphine)palladium(II) dichloride (0.093 g, 0.133 mmol) and sodium carbonate (1.395 g, 13.28 mmol) was added. The reaction was placed under positive argon pressure, sealed, and heated to 70 ℃ for 12 hours. The reaction solvent was evaporated, diluted with water, and extracted with 2:1 EtOAc/ether. The organic was collected, dried, and absorbed onto silica and purified with flash chromatography using a hexanes/EtOAc gradient to isolate the title compound, 1g (0.746 g, 59.2%). 1g (1.515 g, 2.424 mmol) was dissolved in dioxane (24.24 mL) and degassed with argon.

Following 4,4,4',4',5,5,5',5'-octamethyl-2,2'-bi(1,3,2-dioxaborolane) (0.923 g, 3.64 mmol), potassium acetate (0.714 g, 7.27 mmol), Pd2(dba)3 (0.067g, 0.073 mmol), and PCy3 (0.061 g, 0.218 mmol) was added to the reaction. The reaction was placed under an argon atmosphere and heated to 85 ℃ for 12 hours. Afterwards, the reaction solvent was evaporated and the crude reaction was dissolved in EtAOc. The organic layer was washed with 1:1 brine/water 3x, collected, and dried with MgSO4. The organic layer was condensed to generate 1h, which was used without further purification (1.042 g, 89%). 1h (0.100 g, 0.270 mmol) and 2-(4-iodo-1H-pyrazol-1-yl)-N,N-dimethylethanamine (0.071 mg, 0.268 mmol) were added to a 5 mL microwave vial. 4:1 DMF/Water (3 mL) was added to the vial and the reaction was degassed with argon. Following, sodium carbonate (0.065 g, 0.620 mmol), Pd2(dba)3 (5.67 mg, 6.20 µmol), and PCy3 (5.20 mg, 0.019 mmol) were added to the vial. The reaction was sealed and heated under microwave irridation for 30 minutes at 120 ℃. The solvent was evaporated and the crude reaction was transferred to a silica loading column. The reaction was purified *via* flash chromatography using a DCM/MeOH gradient to isolate the desired compound 1i (0.031 mg, 26.2%). MeOH/NH3 (7.0 M, 5 mL) was used to dissolve 1i (0.031 mg, 0.054 mmol). The reaction was sealed and heated to 100 ℃ for 96 hours. The reaction was purified *via* flash chromatography using a DCM/MeOH gradient to isolate the desired compound 2 (0.016 mg, 53.5%).

***Synthesis of Compound 3 (CMP3), Compound 3a (CMP3a) and Compound (CMP3b).***

Compounds 3, 3a, and 3b were synthesized in a similar fashion to that of compound 2 (Fig. S7A). Methyl 2-mercaptoacetate (4.31 mL, 47.1 mmol) was added to MeOH (94 mL) and cooled in an ice bath. Following, sodium hydride (3.39 g, 85 mmol) was slowly added and the reaction was stirred for 1 hour at room temperature. The reaction was cooled to -10 ℃ and ethyl propiolate (5.01 mL, 48.0 mmol) was slowly added. The reaction was stirred at 0 ℃ for 1 hour, room temperature for 1 hour, and heated to 45 ℃ for three hours. Following, the reaction was heated for 12 hours at 35 ℃. All reaction solvent was evaporated and the reaction was redissolved in water, neutralized to pH 7, and extracted 10 times with 3:1 chloroform/IPA. The organic extracts were combined, evaporated, and dissolved in EtOAc. The EtOAc/reaction mixture was filtered through a pad of silica and washed with EtOAc. The organic was collected and condensed to generate the title compound, 3c (3.66 g, 49.1%). 3d was synthesized according to the synthesis of 1e (89%). CMP3d (0.150 mg, 0.454 mmol) was dissolved in THF (5 mL) and cooled to -20 ℃. Following, trimethyl borate (0.103 mL, 0.908 mmol) was added. LDA (0.908 mL, 1.362 mmol) was added to the reaction dropwise over the course of 10 minutes. After about 30 minutes, all starting material was consumed and the boronic acid was generated. In a separate reaction vessel, 1b (0.454 mmol) was dissolved in 7:3 THF/Water and degassed with argon. Sodium carbonate (0.193 g, 1.816 mmol) and Pd2(dppf)Cl2 (8.30 mg, 0.011 mmol) was added and the reaction was heated to 65 ℃. The boronic acid was added dropwise over the course of 10 minutes and the reaction was heated a reflux for 2 hours. The reaction was absorbed onto silica and purified *via* flash chromatography using a DCM/MeOH gradient. The desired compound was isolated and dried, 3e (0.067 mg, 30.7%).

To a degassed (N2 bubbling) solution of the compound of intermediate 3e（721 mg, 1.5 mmol） in 1, 4-dioxane (15 mL) were added bis(pinacolato)diboron (571 mg, 2.25 mmol), KOAc (736 mg, 7.5 mmol) Pd2(dba)3 (41.2 mg, 0.045 mmol), and PCy3 (37.9 mg, 0.135 mmol). Then mixture was heated at 100℃ for 2 hours. The reaction mixture was filtered over a pad of celite and concentrated under vacuum to obtain the crude product 3f which was used in the next step without further purification.

To a solution of crude 3f (735.4 mg, 1.5 mmol) and 3g（596.4 mg, 2.25 mmol）in DMF (12 mL) and water (3 mL) were added Na2CO3 (795 mg, 7.5 mmol), Pd2(dba)3 (41.2 mg, 0.045 mmol), and PCy3 (37.9 mg, 0.135 mmol). Then the reaction mixture was stirred at 120℃ for 30 minutes under microwave irradiation. The reaction mixture was filtered over a pad of celite and concentrated under vacuum. The residue was purified with column chromatography (2-4% MeOH /DCM) to afford the product 3h (233 mg, 40% over two steps) as a yellow solid.

To a solution of 3h (233 mg, 0.4 mmol) in ammonia solution (7M) in methanol (4 mL) was added ammonia solution (1.5 mL, NH3 25-28%), then the reaction mixture was stirred at 80℃ for 5 days. The solvent was removed under reduced pressure. The residue was purified with column chromatography (2-10% MeOH /DCM) to afford the product (439.6, 68%) as a yellow solid.

Preparative chiral HPLC separation of racemic 3i resulted in pure enantiomers 3a and 3b. (Chiralpak AD, EtOH/ACN/DEA=90/10/0.1 (V/V/V), 25ml/min) (Fig. S7B-7D).

CMP3a: 1H NMR (400 MHz, CDCl3) δ 8.13 (d, J = 7.1 Hz, 1H), 7.77 (d, J = 4.1 Hz, 2H), 7.61 (dd, J = 9.8, 7.0 Hz, 4H), 7.51 (t, J = 7.3 Hz, 1H), 7.34 (t, J = 7.3 Hz, 1H), 7.16 (s, 1H), 7.03 (s, 1H), 6.94 – 6.83 (m, 1H), 6.66 (s, 1H), 5.79 (d, J = 5.8 Hz, 1H), 4.31 – 3.96 (m, 2H), 2.89 – 2.51 (m, 2H), 2.21 (s, 6H), 1.70 (d, J = 6.0 Hz, 3H). 13C NMR (101 MHz, CDCl3) δ 163.81, 154.64, 147.62, 140.54, 136.80, 134.63, 133.80, 133.30, 130.29, 128.46, 127.29, 126.50, 126.46 (q, J = 30.2 Hz), 126.11 (q, J = 5.7Hz), 124.38 (q, J = 272.1 Hz), 123.82, 120.43, 119.07, 115.41, 112.97, 112.43, 112.30, 76.26, 58.98, 50.62, 45.56), 24.89. 19F NMR (376 MHz, CDCl3) δ -58.31 (s). HRMS (ESI) m/z calcd for C28H28N6O2SF3 (M+H) + 569.1947, found 569.1922.

***Computational Modeling***

Computational modeling studies were completed using AutoDock Vina, AutoDock Tools, and Discovery Studio 3.5. Using AutoDock Tools, kinase crystal structures were prepared as follows: 1) All hydrogens were added as 'Polar Only'; 2) A grid box for the ATP binding site was created. Compounds to be computationally modeled were assigned appropriate rotatable bonds using AutoDock Tools. To computational model the compounds, AutoDock Vina was employed. AutoDock Vina provides docking scores in terms of ΔG values. After the modeling study, kinase targets with high affinity ΔG values were visualized and analyzed with Discovery Studio 3.5.

*NEK2 Biochemical Inhibition Assay*

Kinase activity was measured in a microfluidics assay that monitors the separation of a phosphorylated product from substrate. The assay was run using a 12-sipper chip on a Caliper EZ Reader II (PerkinElmer®, Walthman, USA) with separation buffer (100 mM HEPES, 10 mM EDTA, 0.015% Brij-35, 0.1% CR-3 [PerkinElmer®, Walthman, USA]). In 96-well polypropylene plates (Greiner, Frickenhausen, Germany) compound stocks (20 mM in DMSO) were diluted into kinase buffer (50 mM HEPES, 0.075% Brij-35, 0.1 % Tween 20, 2 mM DTT, 10 mM MgCl2, and 0.02% NaN3) in 12-point ½log dilutions (2 mM–6.32 nM). Afterwards, 1 µL was transferred into a 384-well polypropylene assay plate (Greiner, Frickenhausen, Germany). The NEK2 enzyme (Invitrogen™, Grand Island, USA) was diluted in kinase buffer to a concentration of 2 nM and 5 µL of the enzyme mixture was transferred to the assay plate. The inhibitors/NEK2 enzyme was incubated for 60 minutes with minor shaking. A substrate mix was prepared containing ATP (Ambresco®, Solon, USA) and 5FAM tagged NEK2 peptide (PerkinElmer®, Walthman, USA) dissolved in kinase buffer, and 5 µL of the substrate mix was added to the assay plate. Running concentrations were as follows: ATP (190 µM), peptide (1.5 µM), compound 12-point ½log dilutions (0.2 mM–0.632 nM). For positive control, no inhibitor was added. For negative control, no enzyme was added. For running control, quizartinib was utilized. The plate was run until 10-20% conversion based on the positive control wells. The following separation conditions were utilized: upstream voltage -500V; downstream voltage,

-1900V; chip pressure -0.8. Percent inhibition was measured for each well comparing starting peptide to phosphorylated product peaks relative to the baseline. Dose response curves, spanning the IC50 dose, were generated in GraphPad Prisim 6 and fit to an exponential one-phase decay line and IC50 values were obtained from the half-life value of the curve. IC50 values were generated in duplicate and error was calculated from the standard deviation between values.

*Supplementary Figures*

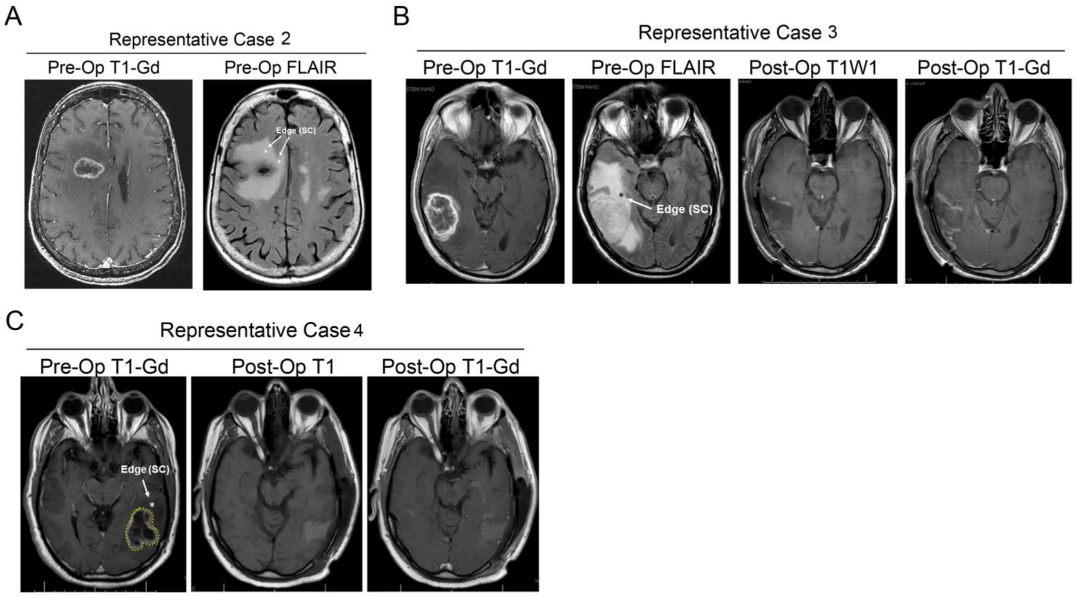


**Fig. S1. MRI-guided intra-operative isolation of glioblastoma core and edge samples.**

(A-C) Representative images to define the tumor core (T1-Gadolinium(+) areas) and edge (T1-Gadolinium(-)/T2-FLAIR abnormal areas in the non-eloquent deep white matter) lesions within the glioblastoma tumors.

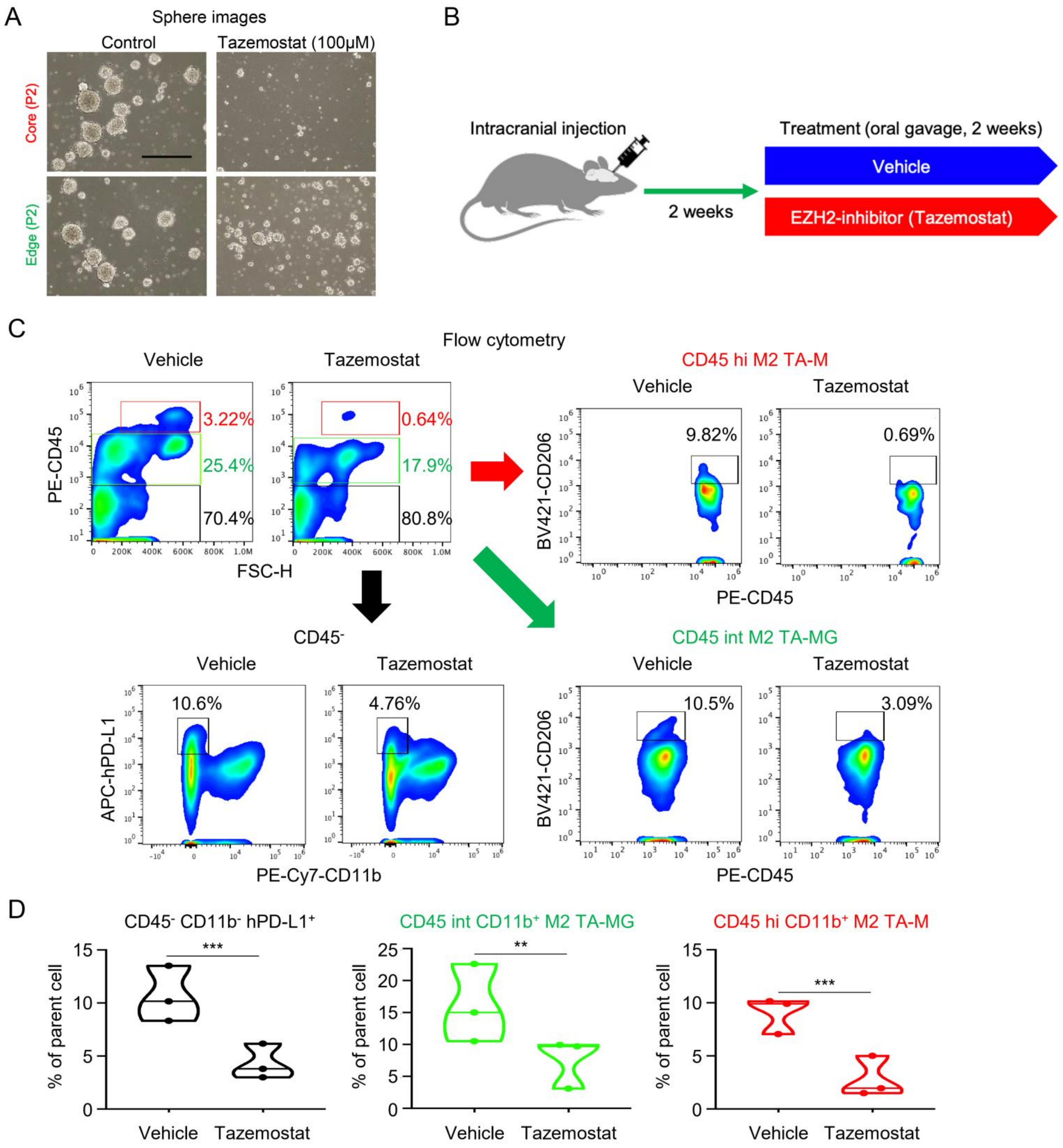


**Fig. S2. Tumor-core derived glioma spheres are more sensitive to EZH2 inhibition.**

(A) Phase bright images of glioma spheres derived from tumor core and tumor edge lesions, followed by Tazemostat treatment at 100μM. The control samples had DMSO treatment. The results indicated that glioma spheres derived from tumor core was more sensitive to Tazemostat compared to its edge counterparts. (B) Scheme for establishment of intracranial mouse tumor models using tumor core-derived glioma spheres, followed by the Tazemostat oral administration. (C) Flow cytometry of the surface CD45 expression of isolated tumor-associated cells pretreated with vehicle (left) or tazemostat (right). Flow cytometry analysis of CD45-CD11b-hPD-L1+ cells (left bottom), CD45 int CD11b + tumor-associated microglia (TAMG; right upper), and CD45 hi CD11b + tumor-associated macrophage (TAM; right bottom). (D) Flow cytometry analysis of the parent cell numbers of total CD45-CD11b-hPD-L1+ cells (left), CD45 int CD11b + TAMG (middle), and CD45 hi CD11b + TAM (right). ** $P < 0.01$ and *** $P < 0.001$, with t test.

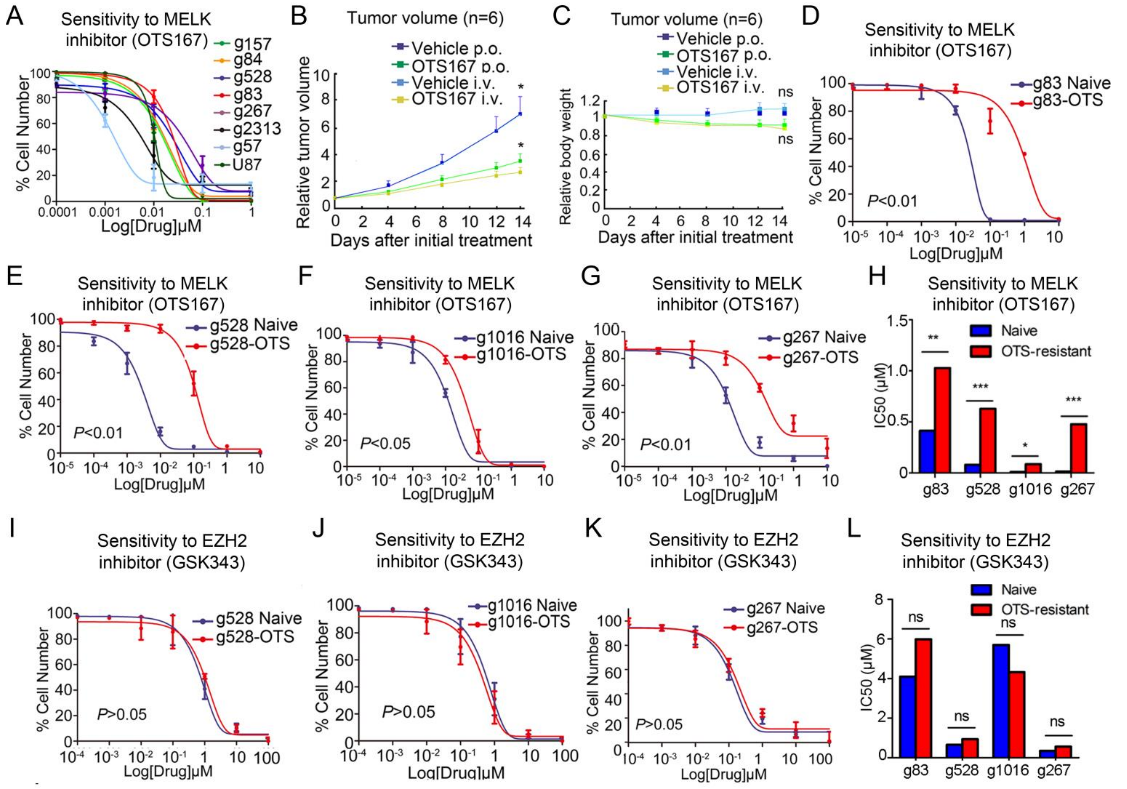


**Fig. S3. NEK2 plays an essential role in post-translational regulation of EZH2.**

(A) Graph showing the relative cell numbers with indicated glioblastoma lines with varying doses of OTS167. OTS167 treatment inhibits cell growth of U87 and the patient-derived glioma sphere lines in a dose-dependent manner. (B) Graph indicating the relative tumor volumes of indicated models. OTS167 treatment attenuates tumor growth in U87 subcutaneous xenograft mouse models (* $P < 0.05$, with one-way ANOVA). (C) Graph indicating no noticeable body weight changes due to OTS167 treatments in U87 subcutaneous xenograft mouse models (ns; not significant. $P > 0.05$, with one-way ANOVA). (D) *In vitro* cell viability assay indicated long-term treatment induced OTS167 resistant population in g83 spheres ($P < 0.01$, with one-way ANOVA). (E) *In vitro* cell viability assay indicated long-term treatment induced OTS167 resistant population in g528 spheres ($P < 0.01$, with one-way ANOVA). (F) *In vitro* cell viability assay indicated long-term treatment induced OTS167 resistant population in g1016 spheres ($P < 0.05$, with one-way ANOVA). (G) *In vitro* cell viability assay indicated long-term treatment induced OTS167 resistant population in g267 spheres ($P < 0.01$, with one-way ANOVA). (H) Graph indicating that IC50s of OTS167 were significantly elevated in resistant population of g83, g528, g1016 and g267 glioma spheres compared to their naïve counterparts (* $P<0.01$, ** $P<0.01$, *** $P < 0.001$, with $t$ test). (I) *In vitro* cell viability assay indicated that OTS resistant g528 spheres exhibited stable sensitivity to EZH2 inhibitor GSK343 ($P > 0.05$, with one-way ANOVA). (J) *In vitro* cell viability assay indicated that OTS resistant g1016 spheres exhibited stable sensitivity to EZH2 inhibitor GSK343 ($P > 0.05$, with one-way ANOVA). (K) *In vitro* cell viability assay indicated that OTS resistant g267 spheres exhibited stable sensitivity to EZH2 inhibitor GSK343($P > 0.05$, with one-way ANOVA). (L) Graph indicating that IC50s of GSK343 remain stable in resistant population of g83, g528, g1016 and g267 glioma spheres (ns $P > 0.05$, with $t$ test).

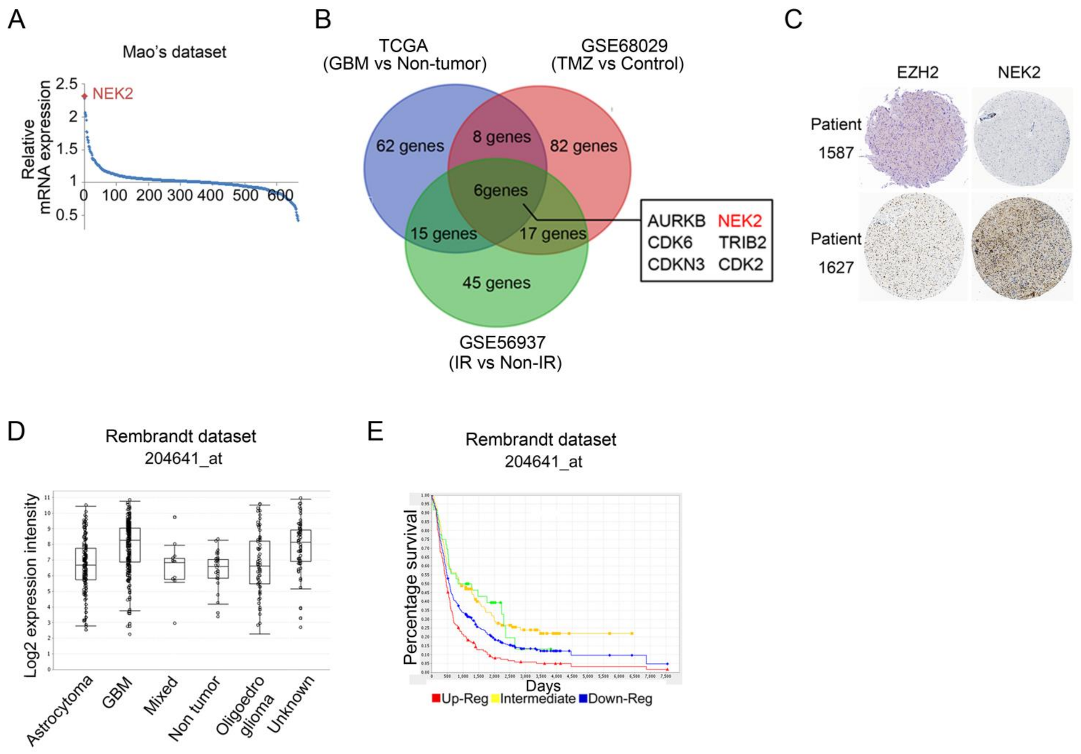


**Fig. S4. NEK2 was correlated with therapy resistance and severe prognosis.**

(A) Graph showing the relative mRNA expression in the glioma sphere lines (n= 6) compared to the normal astrocytes (n= 1), determined by the genome-wide transcriptome microarray analysis (GSE67089). *NEK2* is among the most up-regulated kinases in the glioma spheres. (B) Venn diagram to select for differentially expressed genes indicated that *NEK2* was one of the most differentially expressed kinase-encoding genes correlated to radio resistance, TMZ resistance and glioblastoma subtype. (C) Representative tissue microarray results for EZH2 and NEK2 in glioblastoma samples. NEK2 displayed the strong association with EZH2 expression at protein level in the Human protein Atlas dataset ($EZH2^{High}$ samples n = 69, $EZH2^{Low}$ samples n = 83). (D) Analysis of the Rembrandt database between various glioma grades. *NEK2* mRNA expression was elevated in GBM samples (probe set: 204641_s_at). (E) Analysis of the Rembrandt data for patient survival after separation of these cases into three groups depending on the *NEK2* mRNA levels. The inverted correlation is observed between *NEK2* mRNA expression and post-surgical survival of glioma patients (*P* = 0.0285, *NEK2* Up-Regulated≥ 2.0 Folds, n = 120, *NEK2* Intermediated n = 321, *NEK2* Down-Regulated≥ 2.0 Folds, n = 100, probe set: 204641_s_at).

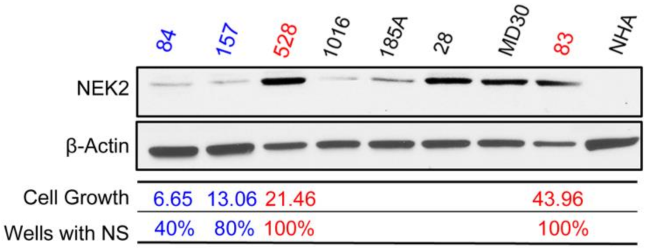


**Fig. S5. NEK2 expression was elevated in glioblastoma cells under TIC-enriching culture condition**

Western blotting analysis showed that NEK2 expression was increased in clonogenic sphere-forming glioma spheres (g528, g83) compared to the spheres exhibiting lower clonogenic ability (g84 and g157). NHA was used as a negative control. β-Actin served as a loading control. Cell growth and sphere formation were observed in 10 days and normalized with day 0.

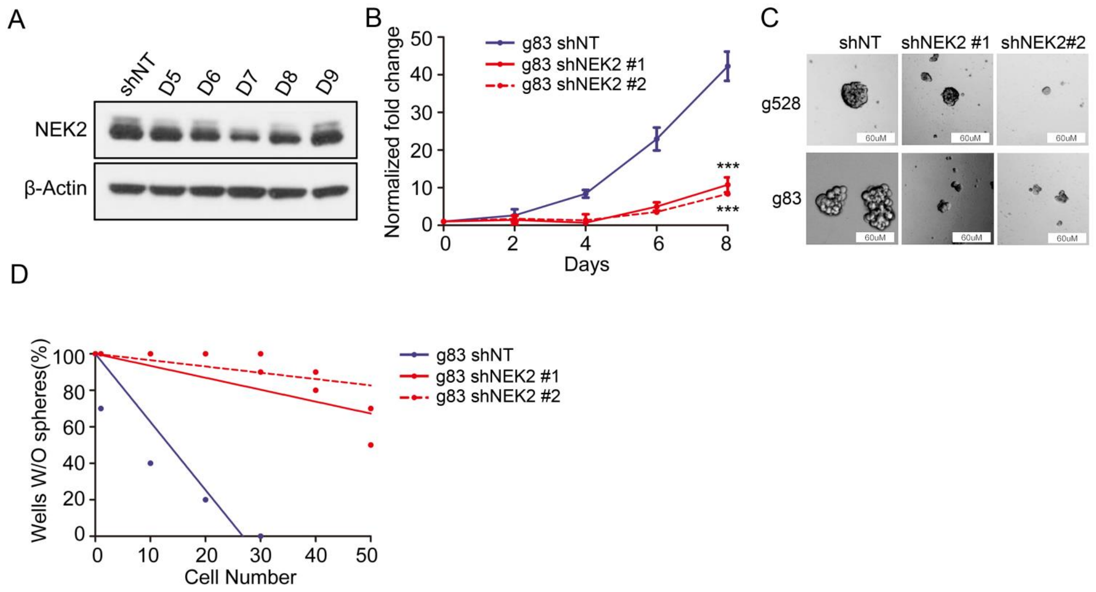


**Fig. S6. Effects of NEK2 knockdown on cell proliferation and self-renewal activity in glioblastoma TICs**

(A) Western blot analysis showed that NEK2 expression was reduced in g83 sphere transduced with shRNA against NEK2 (D5, D6, D7, D8, D9) compared to g83 sphere transduced non-targeting control (shNT). β-Actin served as a loading control. (B) *In vitro* growth assay showing shRNA against NEK2 (shNEK2 #1 and shNEK2 #2) inhibited cell proliferation of g83 spheres (***$P$ < 0.001, n = 6, with one-way ANOVA). (C) *In vitro* clonogenicity was markedly reduced in g528 and g83 spheres transduced with shRNA against NEK2. shNT serves as negative control. (D) *In vitro* clonogenicity assay by limiting dilution neurosphere formation indicated that NEK2 silencing significantly decreased clonogenicity of g83 spheres ($P$ < 0.01, with ELDA analysis).

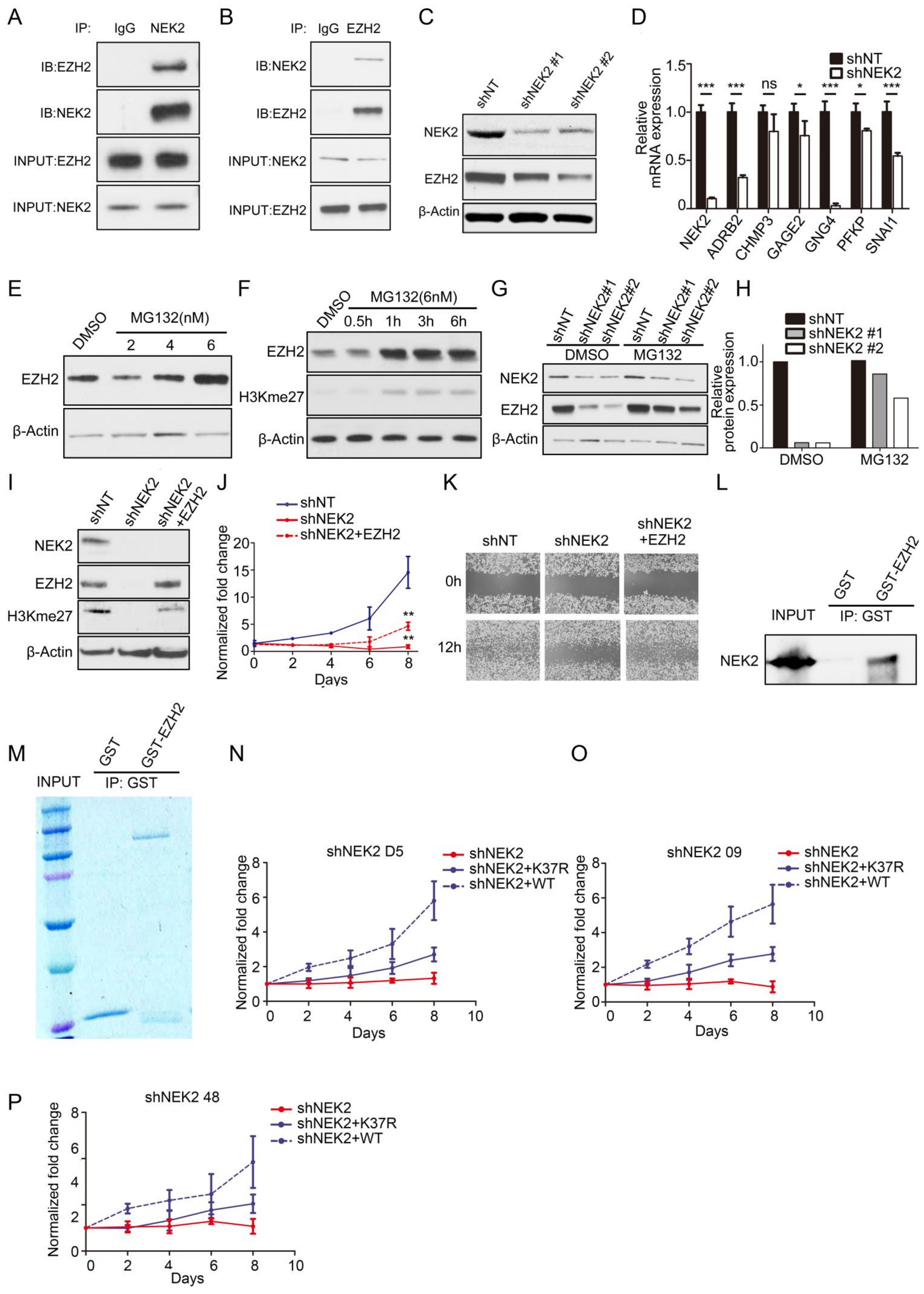


**Fig. S7. NEK2 protected EZH2 from degradation by forming a complex.**

(A) Western blotting analysis of immunoprecipitation using NEK2 antibody or normal mouse IgG showed that NEK2 physically formed a protein complex with

EZH2 in g83 glioma spheres. (B) Western blotting analysis of immunoprecipitation using EZH2 antibody or normal mouse IgG showed that EZH2 physically formed a protein complex with NEK2 in g583 glioma spheres. (C) Western blot analysis indicated that EZH2 expression was significantly reduced in g83 spheres transduced with shRNA against NEK2 (shNEK2 #1 and shNEK2 #2 ). β- Actin served as a control. (D) qRT-PCR analysis showed that expression of multiple downstream targets of EZH2 was reduced by NEK2 KD in g528 spheres (ns $P > 0.05$, * $P < 0.05$, ***$P < 0.001$, n = 3, with *t* test). (E) Western blot analysis for EZH2 was performed in g528 spheres after treated with different dose of MG132 for 6h. The results showed that MG132 blocked the degradation of EZH2 *via* a dose-dependent manner. β-Actin served as a control. (F) Western blot analysis for EZH2 and H3K27me3 in g528 spheres treated with MG132 (6nM). The results showed that MG132 blocked the degradation of EZH2 then increased expression of H3K27me3 *via* a time-dependent manner. β-Actin served as a control. (G-H) Western blotting (G) and quantization analysis (H) of EZH2 in MG132 treated g83 spheres transduced with shRNA against NEK2(shNEK2 #1 and shNEK2 #2) or non-targeting control (shNT ). The results indicated that EZH2 protein underwent degradation after NEK2 silencing and this effect could be reversed by proteasome inhibitor MG13. (I) g267 spheres pretransfected with shRNA against NEK2(shNEK2 #1 and shNEK2 #2) or non-targeting control(shNT) then transduced with EZH2 overexpress vector. Western blot analysis showed that trimethyl-HistoneH3 (Lys27) was reduced by NEK2 KD while could be partially rescued by EZH2 overexpression. β- Actin served as a control.

(J) *In vitro* cell growth assay indicated that NEK2 silencing decreased cell growth of g267 spheres and it could be partially rescued by EZH2 overexpression (**$P < 0.01$, n = 6, with one-way ANOVA). (K) Wound healing assay showed that the ability of cell motility for g267 spheres was decreased by NEK2 silencing and could be rescued by exogenous expression of EZH2. (L-M) The g022 cell lysate and the GST fusion protein are incubated together with glutathione-agarose beads. Complexes recovered from the beads are resolved by SDS-PAGE and analyzed by western blotting, autoradiography or staining (L). The results indicated that NEK2 could be detected in GST-EZH2 immunoprecipitation samples. Coomassie blue staining of GST and GST-EZH2 showed that similar amounts of each protein were used (M). (N-P) *In vitro* cell growth assay indicated that NEK2 silencing *via* 3 3'UTR-shNEK2 lentivirus decreased cell growth of g267 spheres and it could be partially rescued by NEK2-WT overexpression vector but not K37R mutation vector ($P < 0.05$, n = 6, with one-way ANOVA).

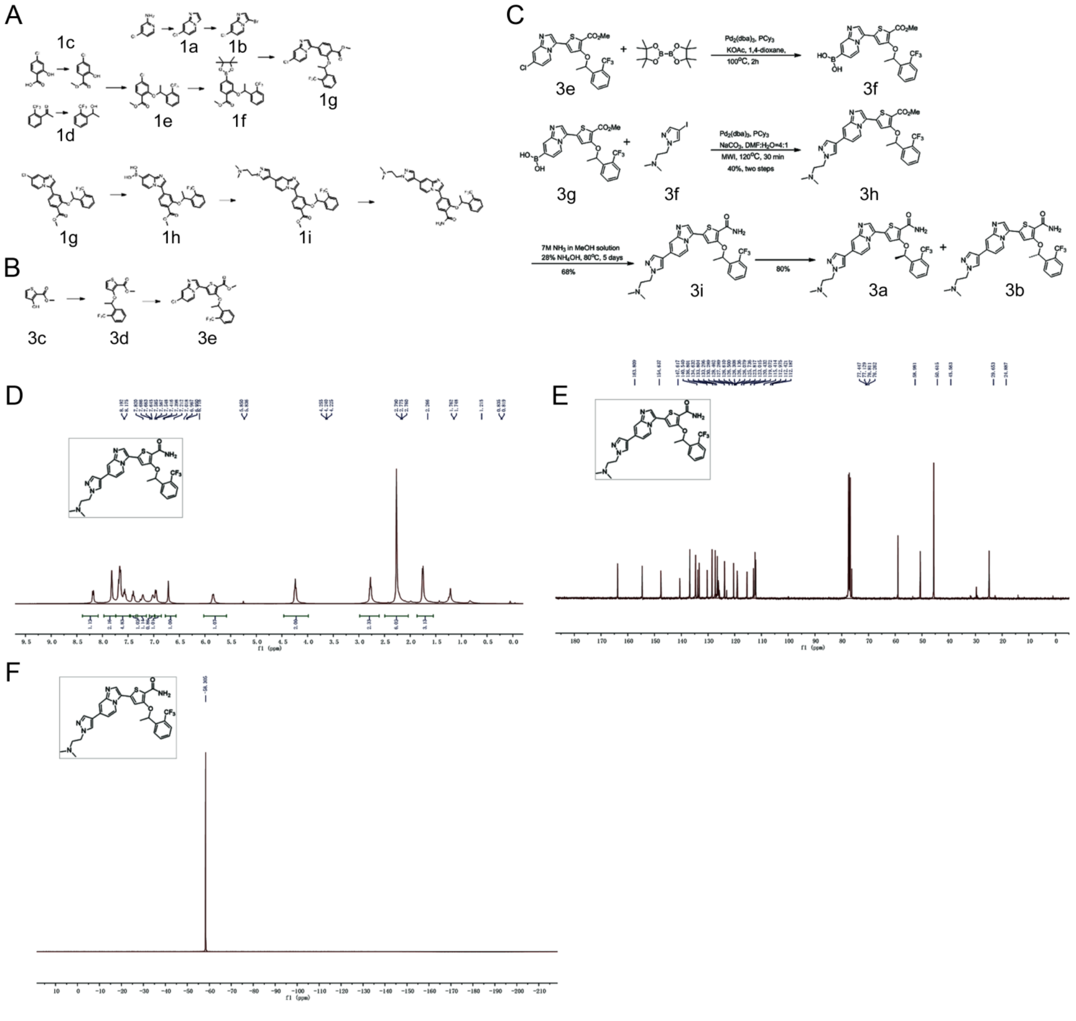


**Fig. S8. Synthesis of NEK2 Inhibitor Compound 3a (CMP3a).**

(A-C) General methods for synthesis of CMP2 (A), CMP3e (B) and CMP3a (C). (D-F) The 1H NMR spectra of compound CMP3a synthesis.

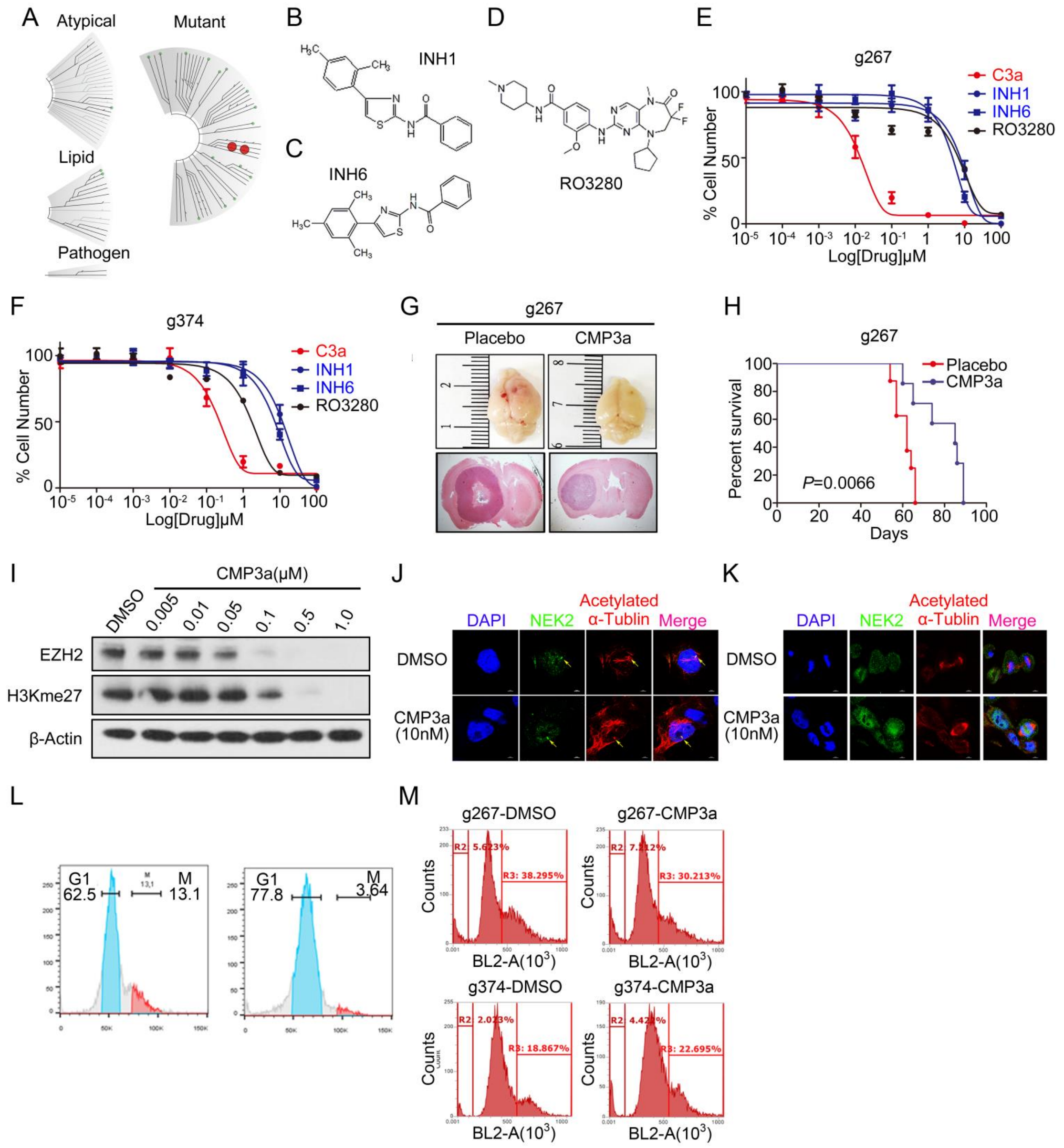


**Fig. S9. NEK2 inhibitor CMP3a increased radio sensitivity and decreased tumor growth in glioblastoma mainly depends on EZH2.**

(A) CMP3a was screened at a 15 nM concentration against mutant kinases representing all kinase clusters by using KINOMEscan. CMP3a showed gratifying specificity to NEK2. (B-D) Chemical structure of INH1, INH6 and RO3280. (E-F) *In vitro* cell viability assay for comparison between CMP3a and other indirect NEK2 inhibitors (INH1 and INH6) in g267 (E) and g374 (F) spheres. The results indicated that CMP3a attenuated cell growth of glioma sphere more efficiently compared with INH1 and INH6. (G) Representative images of mouse brains (upper panel) and H&E stained mouse brain section (lower panel) after the intracranial transplantation of g267 spheres then followed continuously 10-day CMP3a treatment or placebo by tail vein injection. The results showed that CMP3a treatment reduced tumorigenesis of g267 spheres *in vivo*. (H) Kaplan-Meier analysis for mice after the intracranial transplantation of g267 spheres then followed continuously 10-day CMP3a treatment for different does or placebo by tail vein injection. The results showed that CMP3a treatment prolonged the survival of the mice. Data was analyzed with log-rank test. (I) Western blotting analysis showed EZH2 expression was decreased in g267 spheres treated with CMP3a in a dose-dependent manner. β-Actin served as a loading control. (J-K) Immunocytochemistry analysis of NEK2 and acelylated α-Tublin expression in g267 cells treated with CMP3a or DMSO for 12h. DAPI was used for nuclear staining. The results indicated that NEK2 protein was still localized to centrosomes after treatment of CMP3a (J) and the mitosis of glioma sphere cells was not largely affected by CMP3a treatment (K). (L) Flow cytometry data showed that NEK2 was expressed in all phases of cell cycle but not only during division phase in glioblastoma cells. (M) Flow cytometry data showed that inhibition does by CMP3a didn’t affect cell cycle progression of glioblastoma cells.

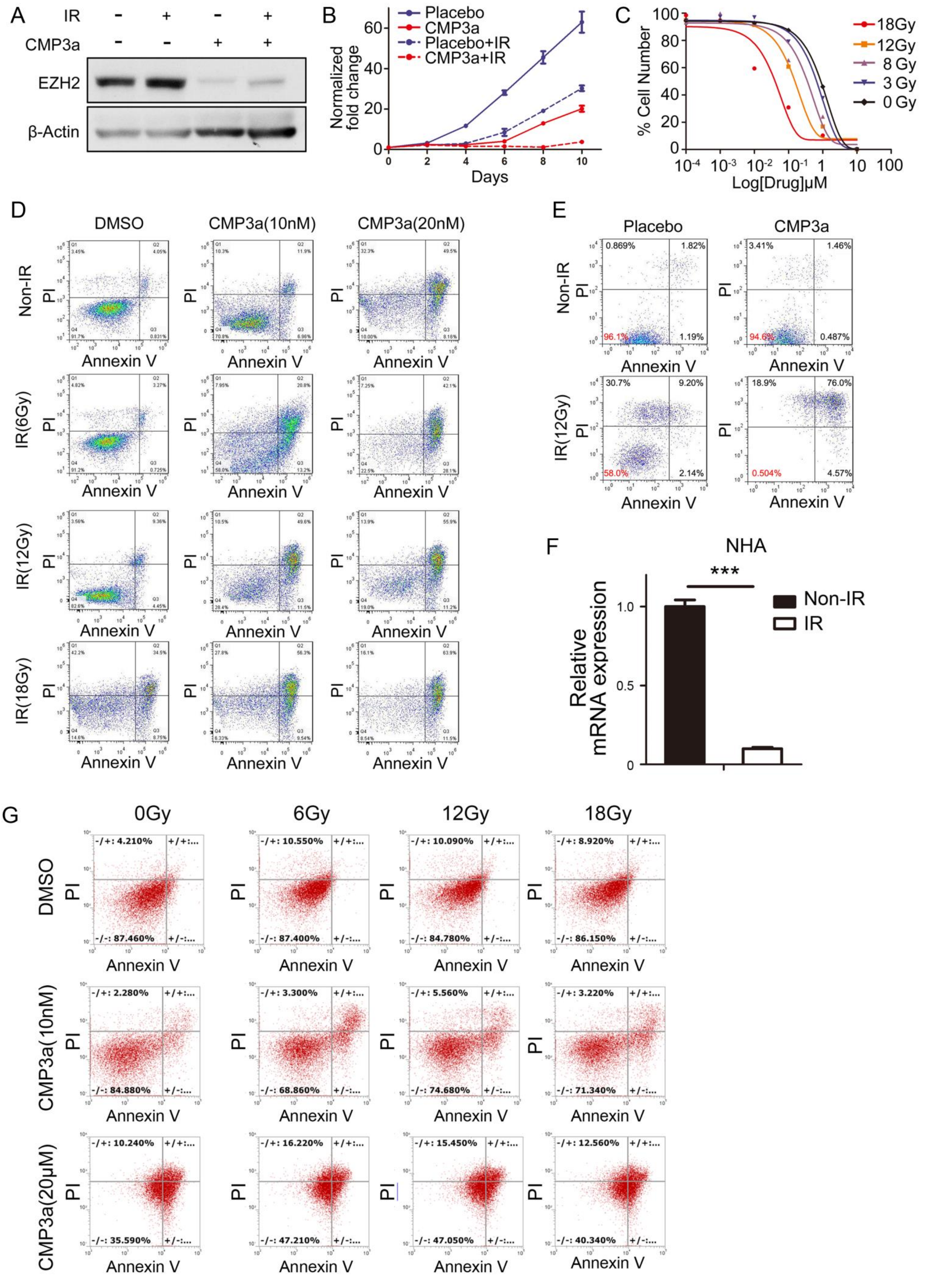


**Fig. S10. NEK2 inhibitor increased radio sensitivity in glioblastoma without affecting normal brain cells.**

(A) Western blot indicated that EZH2 expression was increased after irradiation (12Gy) while could be significantly reduced by CMP3a treatment in g528 spheres. (B) *In vitro* growth assay showed CMP3a inhibited cell proliferation of g528 spheres ($P < 0.001$, n = 6, with one-way ANOVA). (C) *In vitro* cell viability assay indicated that CMP3a treatment combined with irradiation (12Gy) inhibited cell proliferation of g528 spheres in a dose-dependent manner. (D-E) Flow cytometry analysis for apoptosis with Annexin V antibody and Propidium Iodide using g267 (D) or g528 (E) spheres pretreated with CMP3a with or without radiation treatment. The results indicated that CMP3a treatment increased cell apoptosis of g267 or g528 spheres in a dose-dependent manner when combined with irradiation. (F) qRT-PCR data showed that *NEK2* expression was dramatically decreased after IR in NHA cells. (G) Flow cytometry analysis for apoptosis showed that CMP3a did not significantly affect sensitivity of NHA cells to irradiation.

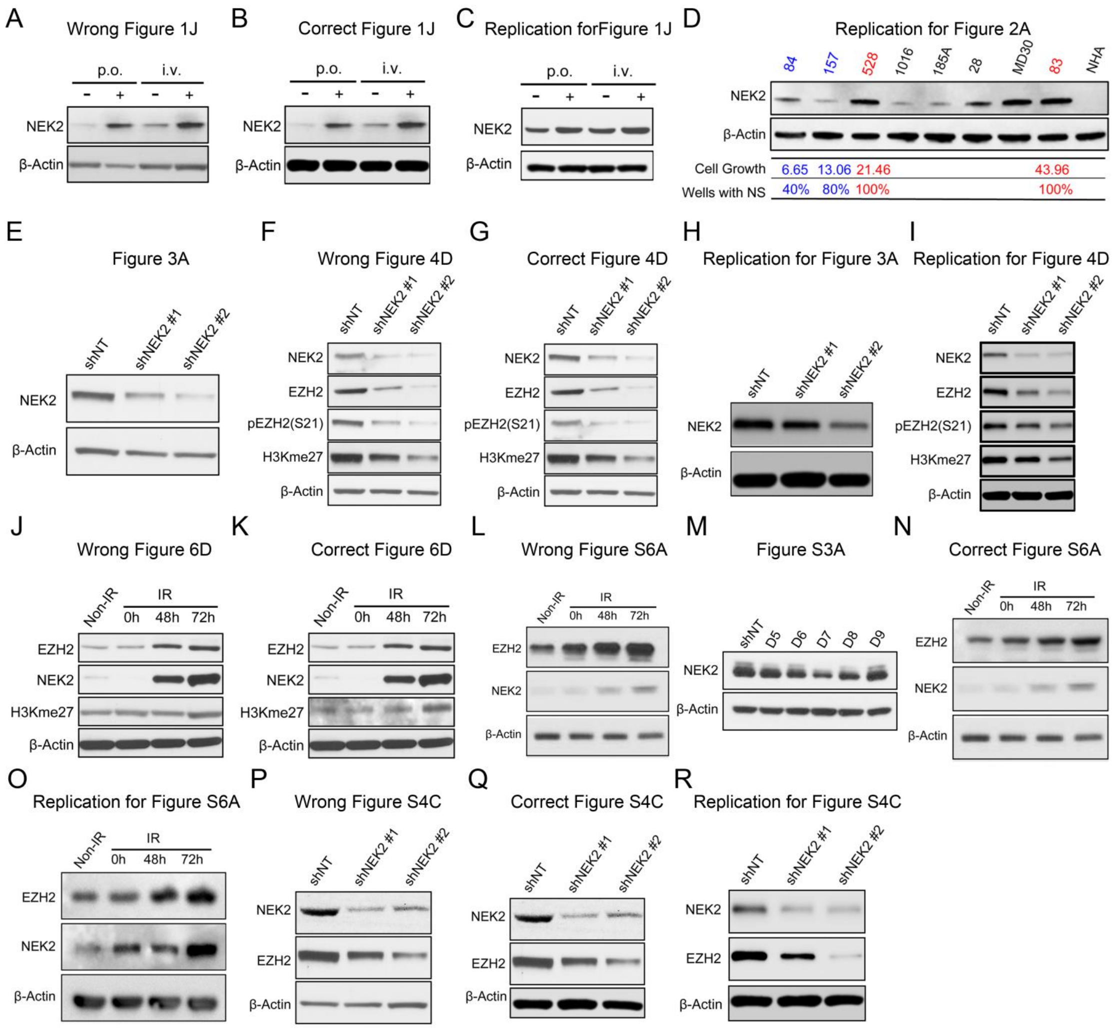


**Fig. S11. Corrections for the wrongly assembled western blotting bands in the original publication.**

(A-D) The actin for Fig. 1J was wrongly assembled (A) with actin from Fig. 2A. We revised Fig. 1J with the correct actin bands (B) and uploaded replicates for Fig. 1J (C) and replicates for Fig. 2A (D) to show that our data was consistent of integrity. (E-I) Fig. 3A (E) and Fig. 4D (F) were using the same samples with NEK2 knock-down in g528 spheres. Fig. 3A (E) was a confirmation for the knock-down efficiency and Fig. 4D (F) was the knock-down effects on the downstream targets of NEK2. We found that Fig. 4D was mislabeled of NEK2 and p-EZH2 (Ser21) and the NEK2 bands should be the same one in these Fig. 3A and Fig. 4D. We revised Fig. 4D (G) and also we submitted replicates for Fig. 3A (H) and replicates for Fig. 4D (I) to show that the results are consistent. (J-K) The H3KMe27 in Fig. 6D was wrongly assembled (J). We then found the correct membrane for H3KMe27 in Fig. 6D and reanalyzed the result. As we can see, similar results could be found with the re-analyzed images. Therefore, we revised Fig. 6D with the correct bands (K). (L-O) The NEK2 band for Fig. S6A (L) was wrongly assembled from Fig. S3A (M). We revised Fig. S6A with the correct actin bands (N). Also we uploaded another replication for Fig. S6A which showed similar trend (O). (P-R) The actin for Fig. S4C (P) was wrongly assembled with the unpaired bands from Fig. 3A (E). We revised Fig. S4C with the correct actin bands (Q) and also attached a replication for Fig. S4C which exhibited the same results (R).

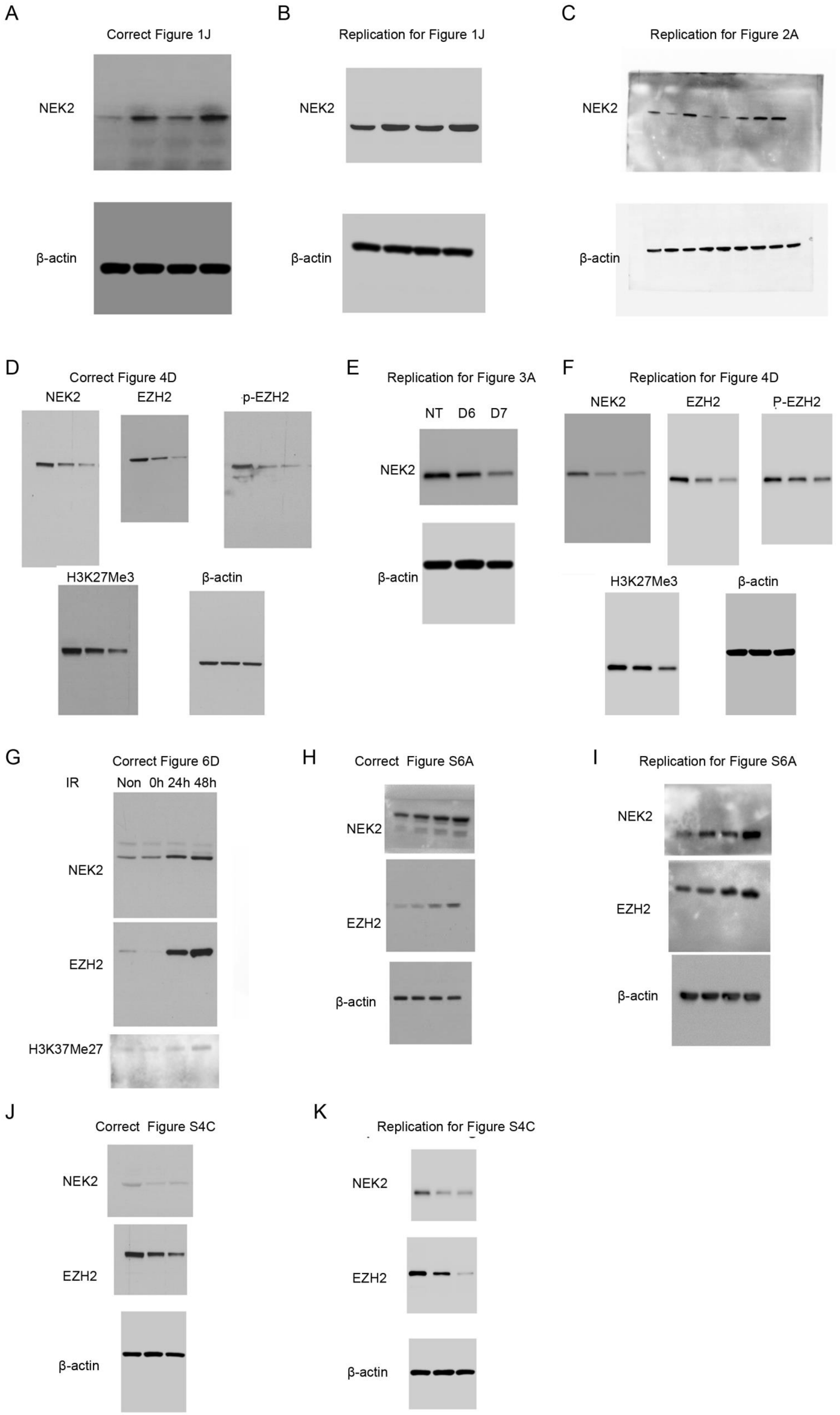
A
Correct Figure 1J
NEK2
β-actin
B
Replication for Figure 1J
NEK2
β-actin
C
Replication for Figure 2A
NEK2
β-actin
D
Correct Figure 4D
NEK2
EZH2
p-EZH2
H3K27Me3
β-actin
E
Replication for Figure 3A
NT D6 D7
NEK2
β-actin
F
Replication for Figure 4D
NEK2
EZH2
P-EZH2
H3K27Me3
β-actin
G
Correct Figure 6D
IR Non 0h 24h 48h
NEK2
EZH2
H3K37Me27
H
Correct Figure S6A
NEK2
EZH2
β-actin
I
Replication for Figure S6A
NEK2
EZH2
β-actin
J
Correct Figure S4C
NEK2
EZH2
β-actin
K
Replication for Figure S4C
NEK2
EZH2
β-actin

**Fig. S12. Uncropped gels for the correction and replication western blotting bands.**

(A) Uncropped gels of correct Fig. 1J. (B) Uncropped gels of replicates for Fig. 1J. (C) Uncropped gels of replicates for Fig. 2A. (D) Uncropped gels of correct Fig. 4D. (E) Uncropped gels of replicates for Fig. 3A. (F) Uncropped gels of replicates for Fig. 4D. (G) Uncropped gels of correct Fig. 6D. (H) Uncropped gels of correct Fig. S6A. (I) Uncropped gels of replicates for Fig. S6A. (J) Uncropped gels of correct Fig. S4C. (K) Uncropped gels of replicates for Fig. S4C.

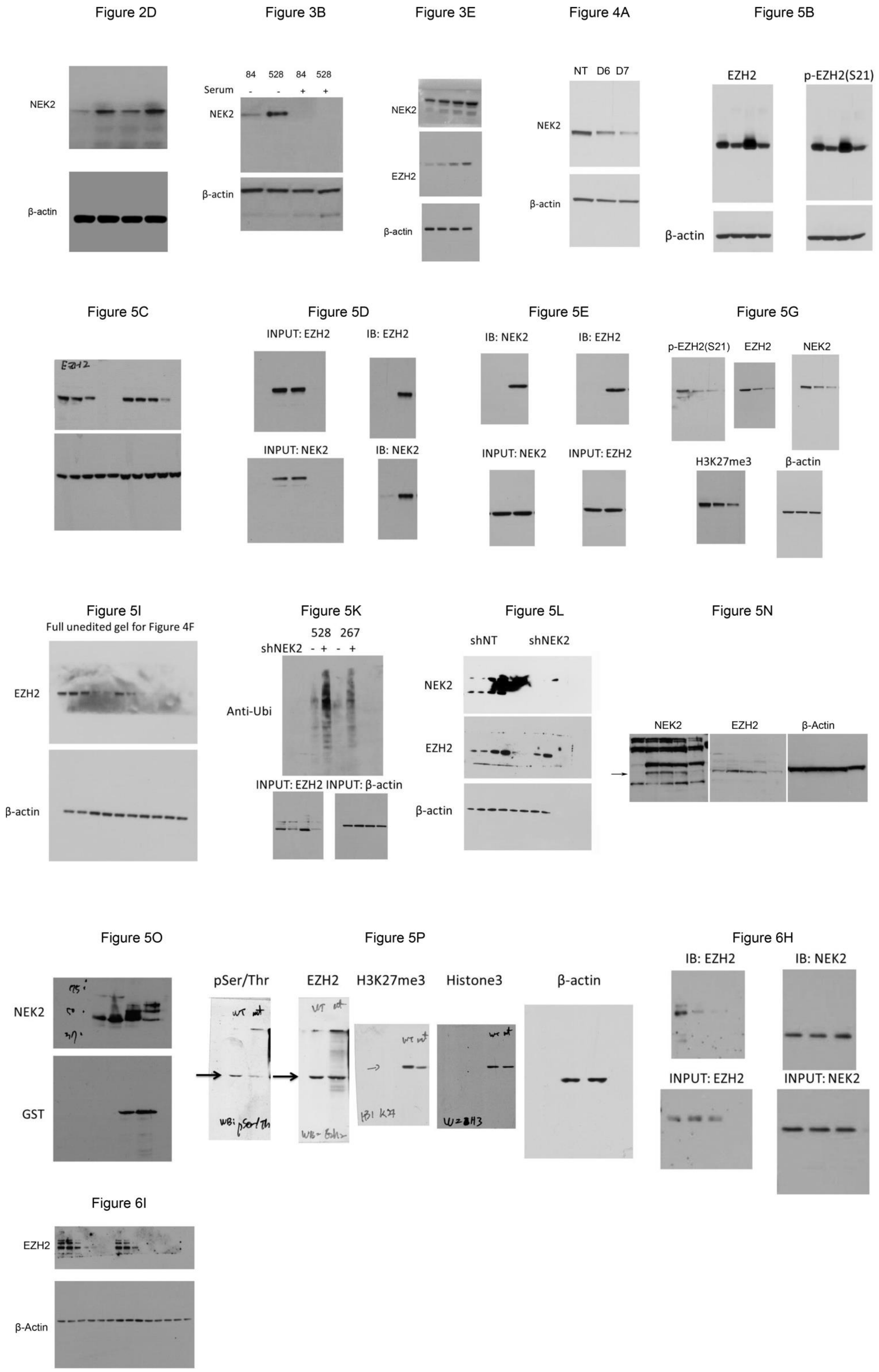


**Fig. S13. Uncropped gels for the western blotting bands in the figures.**

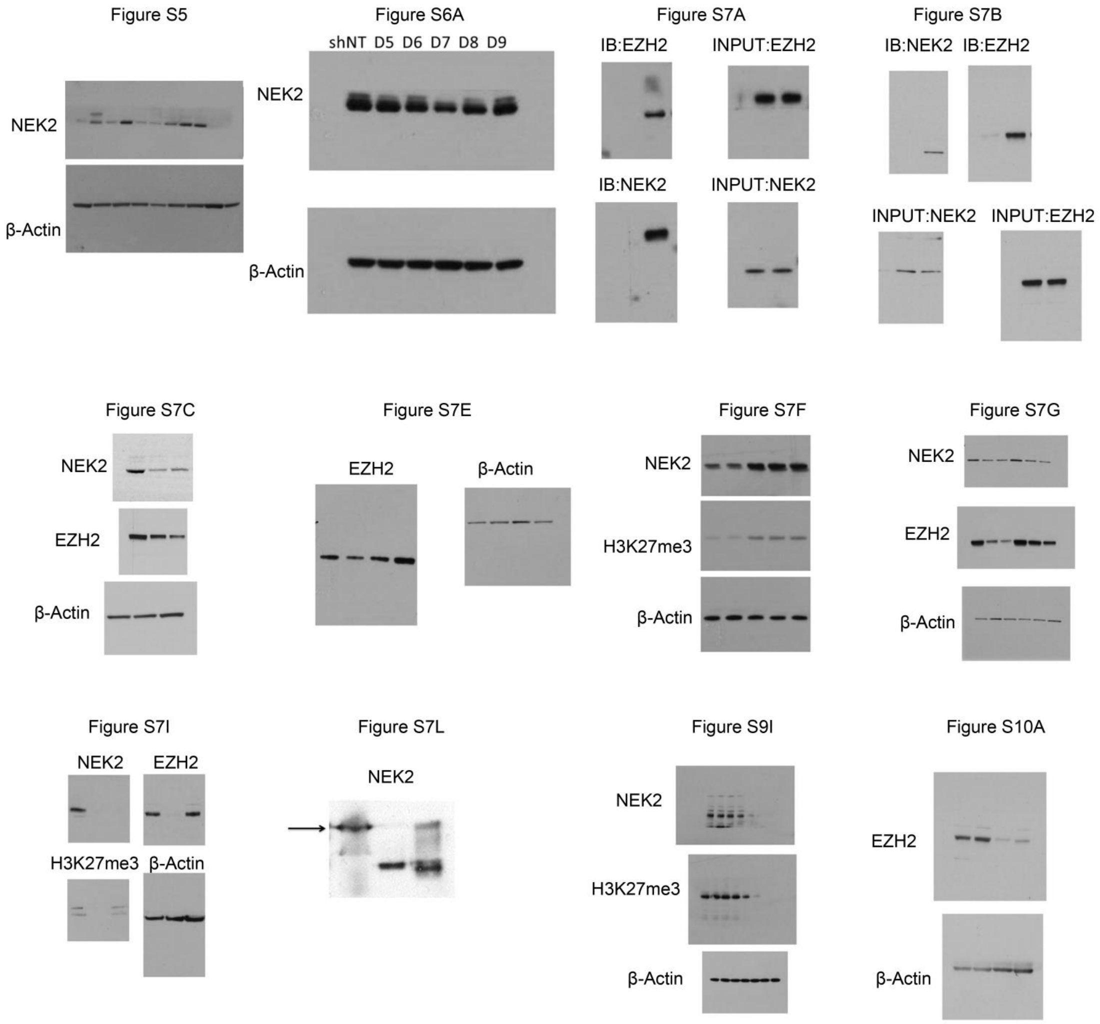


**Fig. S14. Uncropped gels for the western blotting bands in the supplementary figures.**

*Supplementary References*


1. M. E. Ritchie, B. Phipson, D. Wu, Y. Hu, C. W. Law, W. Shi, G. K. Smyth, limma powers differential expression analyses for RNA-sequencing and microarray studies. *Nucleic Acids Res* **43**, e47 (2015).
2. H. Guvenc, M. S. Pavlyukov, K. Joshi, H. Kurt, Y. K. Banasavadi-Siddegowda, P. Mao, C. Hong, R. Yamada, C. H. Kwon, D. Bhasin, S. Chettiar, G. Kitange, I. H. Park, J. N. Sarkaria, C. Li, M. I. Shakhparonov, I. Nakano, Impairment of glioma stem cell survival and growth by a novel inhibitor for Survivin-Ran protein complex. *Clin Cancer Res* **19**, 631-642 (2013).
3. T. Miyazaki, Y. Pan, K. Joshi, D. Purohit, B. Hu, H. Demir, S. Mazumder, S. Okabe, T. Yamori, M. Viapiano, K. Shin-ya, H. Seimiya, I. Nakano, Telomestatin impairs glioma stem cell survival and growth through the disruption of telomeric G-quadruplex and inhibition of the proto-oncogene, c-Myb. *Clin Cancer Res* **18**, 1268-1280 (2012).
4. M. S. Pavlyukov, H. Yu, S. Bastola, M. Minata, V. O. Shender, Y. Lee, S. Zhang, J. Wang, S. Komarova, J. Wang, S. Yamaguchi, H. A. Alsheikh, J. Shi, D. Chen, A. Mohyeldin, S. H. Kim, Y. J. Shin, K. Anufrieva, E. G. Evtushenko, N. V. Antipova, G. P. Arapidi, V. Govorun, N. B. Pestov, M. I. Shakhparonov, L. J. Lee, D. H. Nam, I. Nakano, Apoptotic Cell-Derived Extracellular Vesicles Promote Malignancy of Glioblastoma Via Intercellular Transfer of Splicing Factors. *Cancer cell* **34**, 119-135 e110 (2018).
