## Supplemental Table 1 for "Spatiotemporal Dynamics of Intra-tumoral Dependence on NEK2-EZH2 Signaling in Glioblastoma Cancer Progression"

**Table S1. Sequences of NEK2 shRNAs**

| Name | Sequence |
| --- | --- |
| D5 | CCGGGCCATGCCTTTCTGTATAGTACTCGAGTACTATACAGAAAGGCATGGCTTTTT |
| D6 | CCGGCGTTCGTTACTATGATCGGATCTCGAGATCCGATCATAGTAACGAACGTTTTT |
| D7 | CCGGCGTTACTCTGATGAATTGAATCTCGAGATTCAATTCATCAGAGTAACGTTTTT |
| D8 | CCGGGCAGACGAGCAAAGAAGAAATCTCGAGATTTCTTCTTTGCTCGTCTGCTTTTT |
| D9 | CCGGCCTGTATTGAGTGAGCTGAAACTCGAGTTTCAGCTCACTCAATACAGGTTTTT |
| 3’UTR 09 | CCGGGCCATGCCTTTCTGTATAGTACTCGAGTACTATACAGAAAGGCATGGCTTTTTG |
| 3’UTR 48 | CCGGGCCATGCCTTTCTGTATAGTACTCGAGTACTATACAGAAAGGCATGGCTTTTT |
