## Supplemental Table 2 for "Spatiotemporal Dynamics of Intra-tumoral Dependence on NEK2-EZH2 Signaling in Glioblastoma Cancer Progression"

**Table S2. Primers for qRT-PCR**

| Primer Name | Primer Sequence |
| --- | --- |
| NEK2-For | TCCCCACTGAAATGAACTTTCT |
| NEK2-Rev | CAGCTTGCTAAAGGAACGGA |
| EZH2-For | CCCTTCTCAGATTTCTTCCCA |
| EZH2-Rev | GGACTCAGAAGGCAGTGGAG |
| CD133-For | ACTCCCATAAAGCTGGACCCC |
| CD133-Rev | TCAATTTTGGATTCATATGCCTT |
| ALDH1A3-For | CACTTCTGTGTATTCGGCCA |
| ALDH1A3-Rev | TGGATCAACTGCTACAACGC |
| GAGE2-For | TGGCTATGAGCTTCAGGCTT |
| GAGE2-Rev | AAGAAGGGGAACCAGCAACT |
| STMN3-For | TGCTGTCGCTCATCTGCTCC |
| STMN3-Rev | TCACCTCCATGTCCCCGTACT |
| ADRB2-For | TCCTGGATCACATGCACAAT |
| ADRB2-Rev | GAGCACAAAGCCCTCAAGAC |
| RUNX3-For | GTCTGGTCCTCCAGCTTCTG |
| RUNX3-Rev | CTGTGTTCACCAACCCCAC |
| WNT-For | GGAGGAGGCTACGTTCACAA |
| WNT-Rev | TTTCTGCTACGCTGCTGCT |
| PFKP-For | TGGAGACACTCTCCCAGTCG |
| PFKP-Rev | GGGCCAAGGTGTACTTCATC |
| SNAI1-For | TCTGAGTGGGTCTGGAGGTG |
| SNAI1-Rev | CTCTAGGCCCTGGCTGCTAC |
| CHMP3-For | CCTTCTTGGCAGCATCTTTC |
| CHMP3-Rev | CTGTTTGGAAAGACCCAGGA |
| GNG4-For | CTGGCTTGGGAGATGCTAGT |
| GNG4-Rev | CTTCGCCGGGTTAGTGG |
| GAPDH-For | GAAGGTGAAGGTCGGAGTCA |
| GAPDH-Rev | TTGAGGTCAATGAAGGGGTC |
| NANOG-For | CCTGTGATTTGTGGGCCTG |
| NANOG-Rev | GACAGTCTC CGTGTGAGGCAT |
| ABCG2-For | TATAGCTCAGATCATTGTCACAGTC |
| ABCG2-Rev | GTTGGTCGTCAGGAAGAAGAG |
| CD90-For | ATACCAGCAGTTCACCCATCCAGT |
| CD90-Rev | ATTTGCTGGTGAAGTTGGTTCGGG |
| NOTCH1-For | GAGGCGTGGCAGACTATGC |
| NOTCH1-Rev | CTTGTACTCCGTCAGCGTGA |
| OLIG2-For | CGGCTTTCCTCTATTTTGGTT |
| OLIG2-Rev | GTTACACGGCAGACGCTACA |
| SOX2-For | GTCATTTGCTGTGGGTGATG |
| SOX2-Rev | AGAAAAACGAGGGAAATGGG |
| CD15-For | GAGGGTAGATTGGGGGAAAC |
| CD15-Rev | CACTGCTCGCTGCCTCTC |
| OCT4-For | CATCGGCCTGTGTATATCCC |
| OCT4-Rev | GAAGGAGAAGCTGGAGCAAA |
| 18S-For | GGCCCTGTAATTGGAATGAGTC |
| 18S-Rev | CCAAGATCCAACTACGAGCTT |
