## Supplemental Table 3 for "Spatiotemporal Dynamics of Intra-tumoral Dependence on NEK2-EZH2 Signaling in Glioblastoma Cancer Progression"

**Table S3. Primers for PCR-mediated mutagenesis**

| Name | Sequence |
| --- | --- |
| P1 | AAAATTCGAACCATGCCTTCCCGGGCTGAGGACTATGA |
| P2 | TATAGGATCCGCTAGCGCATGCCCAGGATCTGTCTG |
| P3 | ATGGCAAGATATTAGTTTGGAGAGAACTTGACTATGGCTCCATGACAGAA |
| P4 | TTCTGTCATGGAGCCATAGTCAAGTTCTCTCCAAACTAATATCTT |
